# Genome-wide circadian rhythm detection methods: systematic evaluations and practical guidelines

**DOI:** 10.1101/2020.04.04.024729

**Authors:** Wenwen Mei, Zhiwen Jiang, Yang Chen, Li Chen, Aziz Sancar, Yuchao Jiang

## Abstract

Circadian rhythms are oscillations of behavior, physiology, and metabolism in many organisms. Recent advancements in omics technology make it possible for genome-wide profiling of circadian rhythms. Here, we conducted a comprehensive analysis of seven existing algorithms commonly used for circadian rhythm detection. Using gold-standard circadian and non-circadian genes, we systematically evaluated the accuracy and reproducibility of the algorithms on empirical datasets generated from various omics platforms under different experimental designs. We also carried out extensive simulation studies to test each algorithm’s robustness to key variables, including sampling patterns, replicates, waveforms, signal-to-noise ratios, uneven samplings, and missing values. Furthermore, we examined the distributions of the nominal *p*-values under the null and raised issues with multiple testing corrections using traditional approaches. With our assessment, we provide method selection guidelines for circadian rhythm detection, which are applicable to different types of high-throughput omics data.

**Key points:**

- Various methods have been developed for circadian rhythm detection on a genome-wide scale using omics technologies, yet there has not been a comprehensive summary and evaluation of all existing methods to date.
- Using gold-standard circadian and non-circadian genes, we systematically evaluated the accuracy and reproducibility of seven existing algorithms for circadian rhythm detection on empirical datasets generated from various omics platforms.
- We carried out extensive simulation studies to test each algorithm’s robustness to key variables, including sampling patterns, replicates, waveforms, signal-to-noise ratios, uneven samplings, and missing values.
- We examined the distributions of the nominal *p*-values under the null and raised issues with multiple testing corrections using the Benjamini-Hochberg procedure due to gene-gene correlation and testing being overly conservative.
- We provide method selection guidelines for circadian rhythm detection, which are applicable to different types of high-throughput omics data.

## BACKGROUND

Circadian rhythms are approximately 24-hour oscillations of behavior, physiology, and metabolism that exist in almost all living organisms ranging from prokaryotes to mammals [1, 2]. Circadian rhythm is regulated by the circadian system, which consists of many “clock-controlled genes” that exhibit oscillatory patterns [1]. These oscillations provide organisms with an adaptive advantage by enabling them to predict and adjust to the variations within their environments [3]. Additionally, and perhaps more importantly, disruptions of circadian rhythms have shown to contribute to numerous diseases, including metabolic disorders, heart disease, and aging [4–7]. It is, therefore, of great importance and interest to perform genome-scale analysis of biological rhythms.

Recent advances in omics technologies, including both microarrays and next-generation sequencing, offer appealing platforms to identify circadian genes on a genome-wide scale. These have, indeed, led to the proposal of multifarious methodologies adopted from various fields including mathematics, statistics, astrophysics, etc. The earliest of the selected methods is Lomb-Scargle (LS) periodogram [8], an algorithm adapted from astrophysics that detects oscillations by comparing the data to sinusoidal reference curves of varying periods and phases [9, 10]. ARSER is an algorithm that employs autoregressive spectral estimation to predict periodicity and applies a harmonic regression model to fit the time-series [11]. Unlike the model-based LS and ARSER, JTK_CYCLE is a non-parametric method that detects oscillations by comparing the ranks of the measured values to a set of prespecified symmetric reference curves [3]. Both RAIN and eJTK_CYCLE build on the strengths of JTK_CYCLE: RAIN includes an additional set of asymmetric waveforms and examines the increasing and decreasing portions of the curve separately [12]; eJTK_CYCLE improves JTK_CYCLE by explicitly calculating the null distribution such that it accounts for multiple hypothesis testing and by including non-sinusoidal reference waveforms [13]. Based on the successes of the aforementioned methods, MetaCycle proposes an ensemble framework that integrates results from three different algorithms, LS, ARSER, and JTK_CYCLE [14]. Specifically, MetaCycle detects periodicity using the best of breed methods: its *p*-values are generated using Fisher’s method; its periods and phase estimations are integrated using arithmetic and circular means; and a new periodic model, formulated from ordinary least squares method, is applied to recalculate the amplitude. The most recent method, BIO_CYCLE, is a deep neural network trained on both simulated and empirical circadian and noncircadian time-series [15]. More general information and characteristics of each method are summarized in Table 1.

**Table 1.**
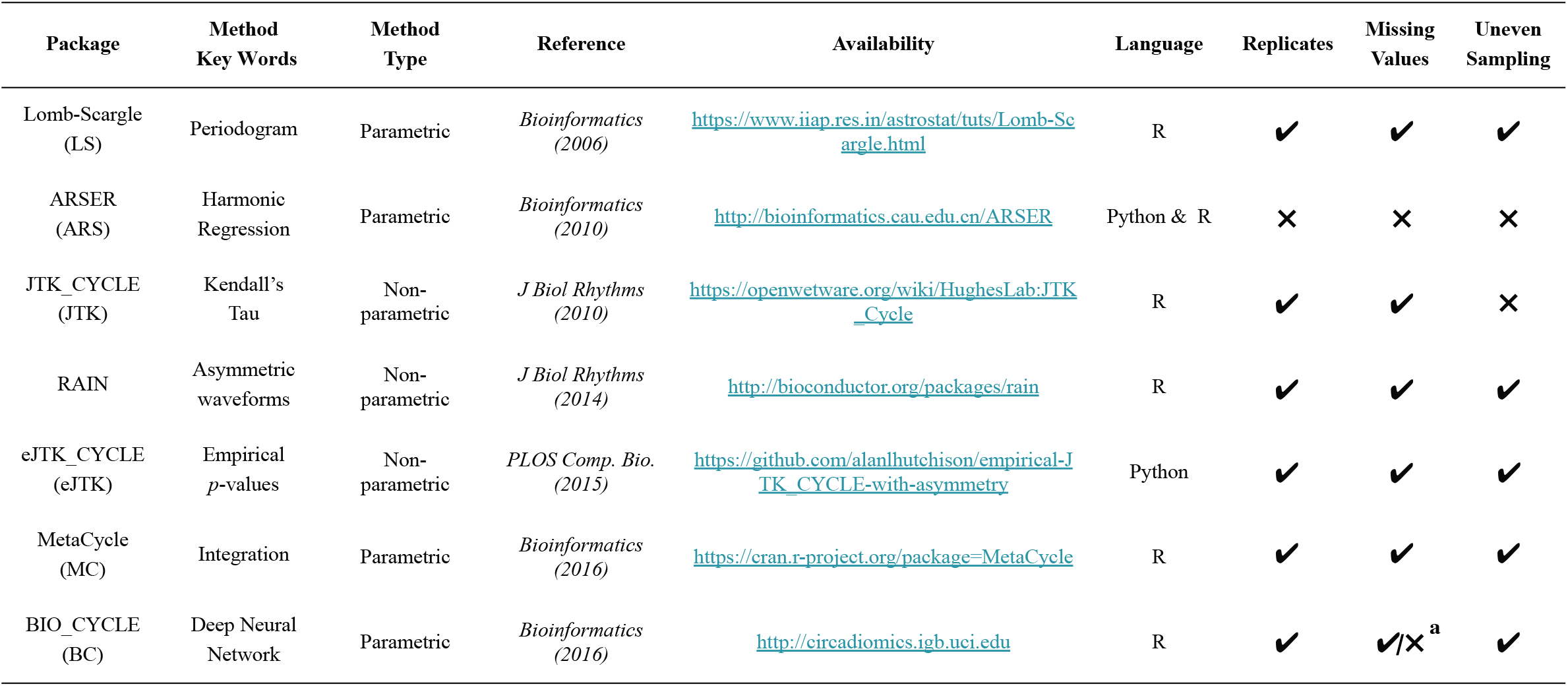
Summary of seven existing methods for circadian rhythm detection.^a^. BIO_CYCLE can be applied to datasets with missing values only if there are replicates and the missingness only pertains to part of the replicates.

Multiple studies [10, 16, 17] have evaluated the performance of different methods for circadian rhythm detection, showing discrepancies among the methods, whose performances depend on multiple factors including experimental designs, waveforms of interest, etc. However, there has not been, to our best knowledge, a comprehensive summary and evaluation of all existing methods to date. Here, we systematically assess the performance of the seven aforementioned algorithms for circadian rhythm detection: LS, ARSER, JTK_CYCLE, RAIN, eJTK_CYCLE, MetaCycle, and BIO_CYCLE.

Specifically, we demonstrated and benchmarked the algorithms using real datasets with gold-standard circadian and non-circadian genes. All empirical data were generated using the liver tissue from *Mus musculus* that had undergone two different experimental designs. Under the dark-dark experimental design (24-hour darkness), we focused on using data from gene expression microarrays to assess the accuracy and reproducibility of each algorithm; under the light-dark experimental design (12-hour light followed by 12-hour darkness), we adopted four different next-generation sequencing platforms and explored the robustness of each method in identifying circadian genes. Furthermore, to extend our assessment to non-transcriptomic datasets, we included a proteomic dataset in our evaluation. In addition, we carried out extensive simulation studies to study how key variables, including sampling patterns, replicates, waveforms, signal-to-noise ratios, uneven samplings, missing values, affect the performance of each method. Lastly, we point out the flaw with using the Benjamini-Hochberg procedure to control for false discovery rate. Through these, we offer guidelines on experimental designs as well as best practices and methods of choice to increase the rigor and reproducibility in the analysis of large-scale circadian rhythms. To assist with the comparison of future methods and datasets using our framework, we provide detailed vignettes on applications of existing methods and performance evaluations with source code available at https://github.com/wenwenm183/Circadian_Genes_Benchmark.

## RESULTS

### Performance assessment using empirical datasets with dark-dark design

We first adopted three gene expression microarray datasets from Hughes et al. [18], Hughes et al. [19], and Zhang et al. [20]. For all three studies, mouse liver samples were collected in every hour or every two-hour under the dark-dark experimental design for 48 hours. We named these three datasets after the first author’s last name and the year of publication as Hughes 2009, Hughes 2012, and Zhang 2014, respectively. In addition, we generated a new downsampled dataset from the Hughes 2009 dataset by keeping the even time-points only, and named it “Downsampled Hughes 2009”. Refer to Table 2A for details of the data. Figure 1 shows the scaled gene expression levels of four known circadian and four non-circadian genes. The circadian genes, including the well-studied *Clock, Cry1, Npas2*, and *Per1* [10], show oscillatory patterns that can be well reproduced across studies, while the non-circadian genes exhibit only noisy signals.

**Table 2.** High-throughput mouse liver datasets adopted for circadian rhythm detection. (A) Dark-dark experimental design. (B) Light-dark experimental design.

| Design | Name | Reference | Accession Number | Tissue Type | Sequencing Platform | Number of Time Points & Replicates | Number of Genes | Time Points |
| --- | --- | --- | --- | --- | --- | --- | --- | --- |
| (A) Dark-Dark | Hughes et al. | *PLOS Genetics* (2009) | GSE11923 | Liver | Microarray | 48 x 1 | 13,029 | CT18, 19, 20, ..., 65 |
|  | Hughes et al. (downsampled) | *PLOS Genetics* (2009) | GSE11923 | Liver | Microarray | 24 x 1 | 12,506 | CT18, 20, 22, ..., 64 |
|  | Hughes et al. | *PLOS Genetics* (2012) | GSE30411 | Liver | Microarray | 24 x 1 | 14,413 | CT0, 2, 4, ..., 46 |
|  | Zhang et al. | *PNAS* (2014) | GSE54652 | Liver | Microarray | 24 x 1 | 20,307 | CT18, 20, 22, ..., 64 |
| (B) Light-Dark | Fang et al. | *Cell* (2014) | GSE59486 | Liver | GRO-seq | 8 x 1 | 17,463 | ZT1, 4, 7, 10, 13, 16, 19, 22 |
|  | Menet et al. | *eLIFE* (2012) | GSE36872 | Liver | Nascent-seq | 6 x 2 | 17,917 | ZT0, 4, 8, 12, 16, 20 |
|  | Menet et al. | *eLIFE* (2012) | GSE36871 | Liver | RNA-seq | 6 x 2 | 17,222 | ZT2, 6, 10, 14, 18, 22 |
|  | Yang et al. | *PNAS* (2018) | GSE109938 | Liver | XR-seq (TS) | 6 x 2 | 17,652 | ZT0, 4, 8, 12, 16, 20 |

**Figure 1.**
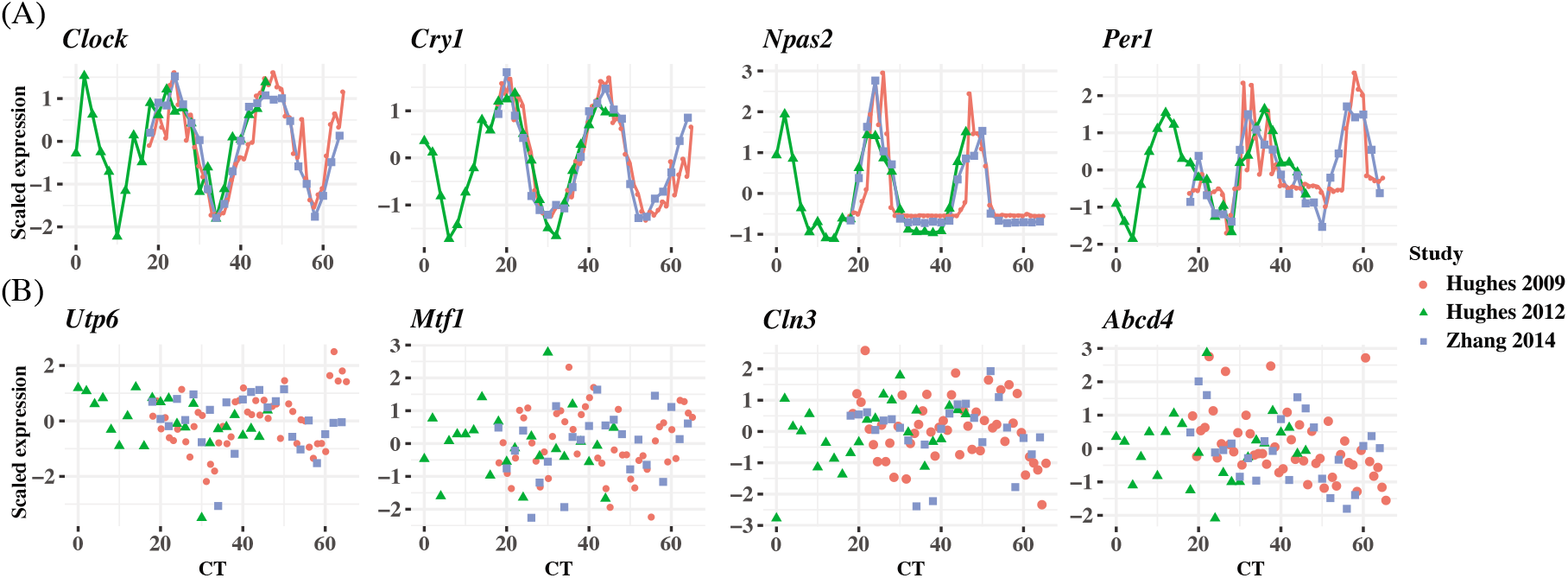
Examples of circadian and non-circadian benchmark gene expressions among three datasets with dark-dark experimental design. Scaled gene expressions from selected (A) circadian genes including *Clock, Cry1, Npas2*, and *Per1* and (B) non-circadian genes including *Utp6, Mtf1, Cln3, Abcd4*.

We set out to apply the seven algorithms to these four datasets to detect significantly cyclic genes and evaluate their performances using 104 circadian [10] and 113 non-circadian genes [21] from previous studies (Supplementary Table 1). The accuracy of each method in Hughes 2009, Downsampled Hughes 2009, Hughes 2012, and Zhang 2014 was first assayed with the precision and recall rates for each algorithm given three *p*-value thresholds, 0.000005 (Bonferroni), 0.00005, 0.0005, and one *q*-value threshold 0.05 (Benjamini-Hochberg). Due to the tradeoff between sensitivity and specificity, with more relaxed thresholds of significance, the precision rates of all methods decrease while the recall rates increase – the 0.05 *q*-value threshold achieves the lowest precision rate yet the highest recall rate for any given method (Figure 2A). While there does not exist a single method that consistently achieves the highest precision or recall rate, JTK_CYCLE and BIO_CYCLE are more effective in controlling for false positives while still detecting true circadian genes. For the other methods, however, there is a much higher variability in precision, especially in the Zhang 2014 dataset (Figure 2A). RAIN and MetaCycle tend to have the highest sensitivity/recall, but this can come with significant sacrifice on precision (Figure 2A).

**Figure 2.**
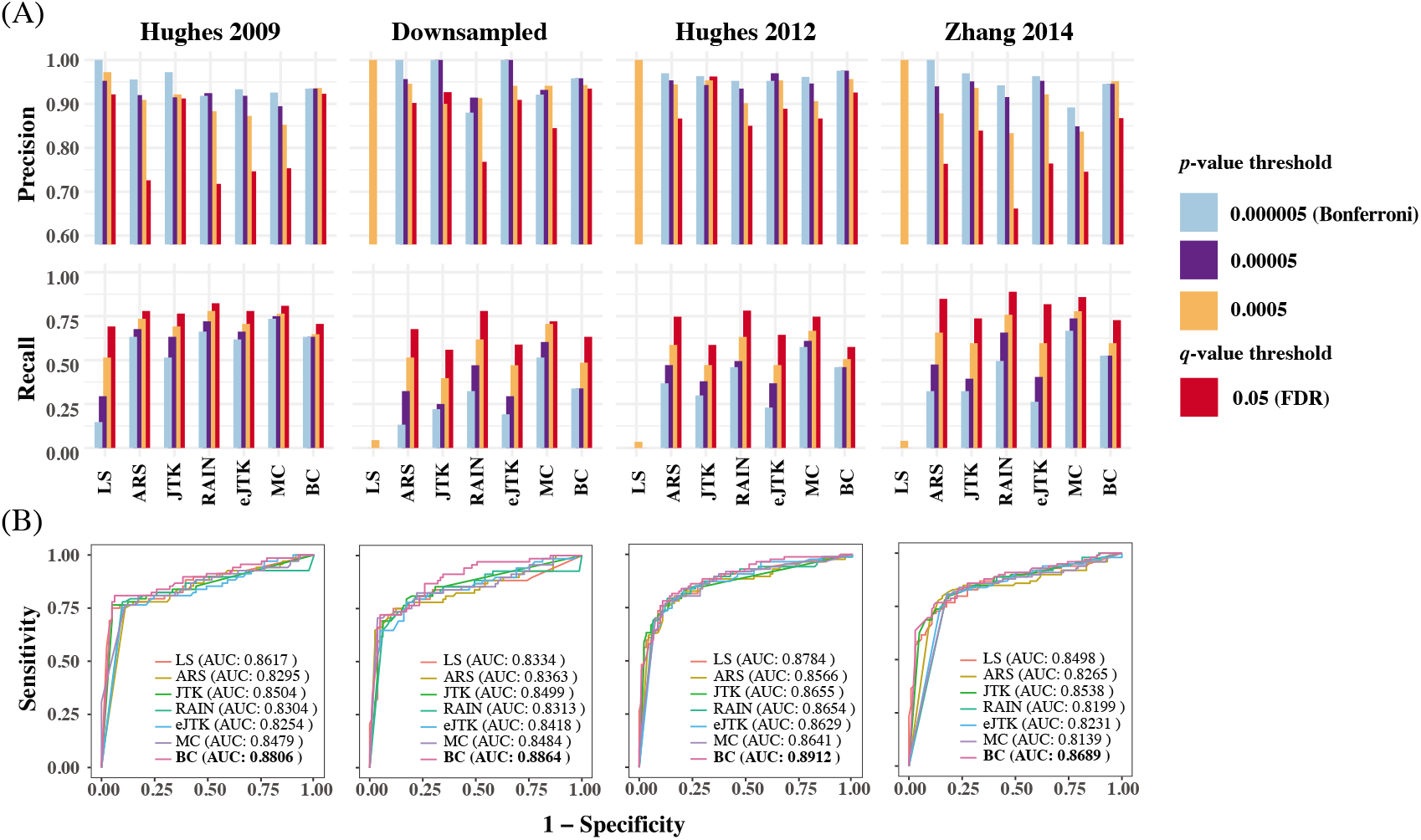
Evaluation of seven methods by precision, recall rates and ROC curves. (A) A *p*-value threshold of 0.000005 (Bonferroni threshold), 0.00005, 0.0005, and a *q*-value threshold of 0.05 (FDR threshold) are adopted for each of the seven methods applied to the four dark-dark empirical datasets. A more relaxed threshold results in a higher recall rate, with FDR being the most sensitive, yet this also leads to a higher number of false positives with a lower precision rate. (B) ROC curves and AUC values using gold-standard circadian and non-circadian genes. Each method is evaluated across four dark-dark empirical datasets. Sensitivity and specificity are calculated using the nominal *p*-values by each method with varying threshold. BIO_CYCLE returns the highest AUC.

In addition, we find that higher sampling frequency can significantly improve the recall rates of all methods. While MetaCycle and RAIN achieve the apparently higher recall rate under different thresholds in dataset sampled at a lower frequency (2 h/2 days), all methods, except for LS, produce comparable recall rates when applied to the Hughes 2009 dataset, which is sampled at 1 h/2 days (Figure 2A). Notably, when analyzing the three datasets with lower sampling frequencies, LS failed under all circumstances with recall rates less than 0.1 (Figure 2A). This is due to the extreme *p*-value distribution of the method with a spike at one, which we will discuss in more detail under “Correlated multiple testing and non-uniform distribution of *p*-values under the null”.

We further computed with the receiver operating characteristic (ROC) curves with a varying threshold on the nominal *p*-values returned by each method (Figure 2B). The area under the curve (AUC) values serve as a joint measure of sensitivity and specificity and are above 0.80 across all benchmark results, suggesting that all methods achieve good sensitivities while controlling for false positive rates. BIO_CYCLE, the deep-learning-based method, achieves the best performance with the highest AUC across all datasets (Figure 2B).

### Reproducibility assessment using empirical datasets with dark-dark design

Reproducibility is one of the core principles for any bioinformatic tools and yet it remains a challenge in the field of circadian rhythm detection, which has not been fully explored. To evaluate the reproducibility of the methods, we first compared and contrasted the significantly cyclic genes returned by each method across the four datasets. To make the input dimensions compatible, we selected a total of 7,570 common genes that are shared across datasets and adopted a *q*-value threshold of 0.05 for significance. The Venn diagrams in Figure 3A show the overlapping relationships of the significant genes returned by each method. While the experimental designs are the same and the observed gene expression measurements are highly concordant (Figure 1), significant discrepancies of the calling results are observed. Of the seven benchmarked methods, ARSER resulted in 721 overlapping significant genes, which is the highest. This is followed by RAIN, eJTK_CYCLE, MetaCycle, BIO_CYCLE, JTK_CYCLE, and LS with 613, 528, 485, 296, 204, and 0 mutually identified positives, respectively. As mentioned previously, LS failed in detecting any significant oscillations for three out of the four datasets.

**Figure 3.**
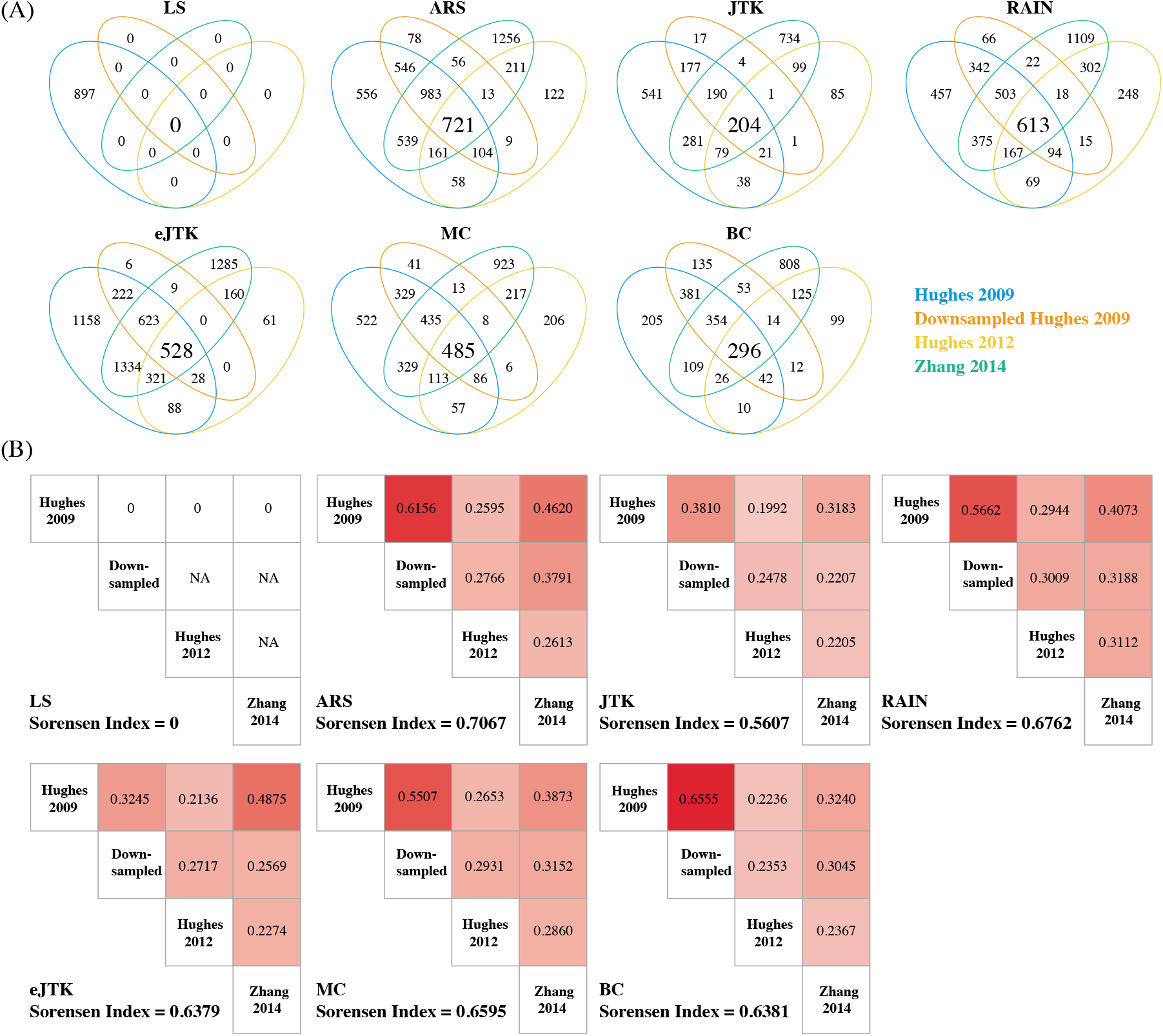
Evaluation of method reproducibility. (A) Venn diagrams display the number of cyclic genes that are significant by each method among the four dark-dark datasets. (B) Jaccard index and the Sorensen index are used as metrics for reproducibility for each method across the four datasets with the same experimental design.

To further assess the reproducibility of the methods, we computed the Jaccard index and the Sorensen index to measure the similarities among the results from each method. Details of these metrics are included in the Materials and Methods section. As a result, RAIN achieves one of the highest Jaccard indices for any pair of comparisons and ARSER achieves the highest overall Sorensen index across all datasets (Figure 3B). On the other hand, our results indicate that JTK_CYCLE, eJTK_CYLE, and BIO_CYCLE produce the lowest similarity metrics across all comparisons (Figure 3B).

### Performance assessment using empirical datasets with light-dark design

Next, we adopted four datasets that underwent light-dark experimental design using different next-generation sequencing platforms (i.e., RNA-seq [22], Nascent-seq [22], GRO-seq [23], and XR-seq [24]) and named each one after its sequencing protocol (Table 2B). The four datasets have much fewer numbers of time-points compared to the datasets from the dark-dark design, yet three of the four datasets have technical replicates (Table 2B). More details of the data can be found in the Materials and Methods section. The oscillatory patterns of known circadian genes are apparent and similar among the various sequencing technologies (Figure 4A), indicating good data quality.

**Figure 4.**
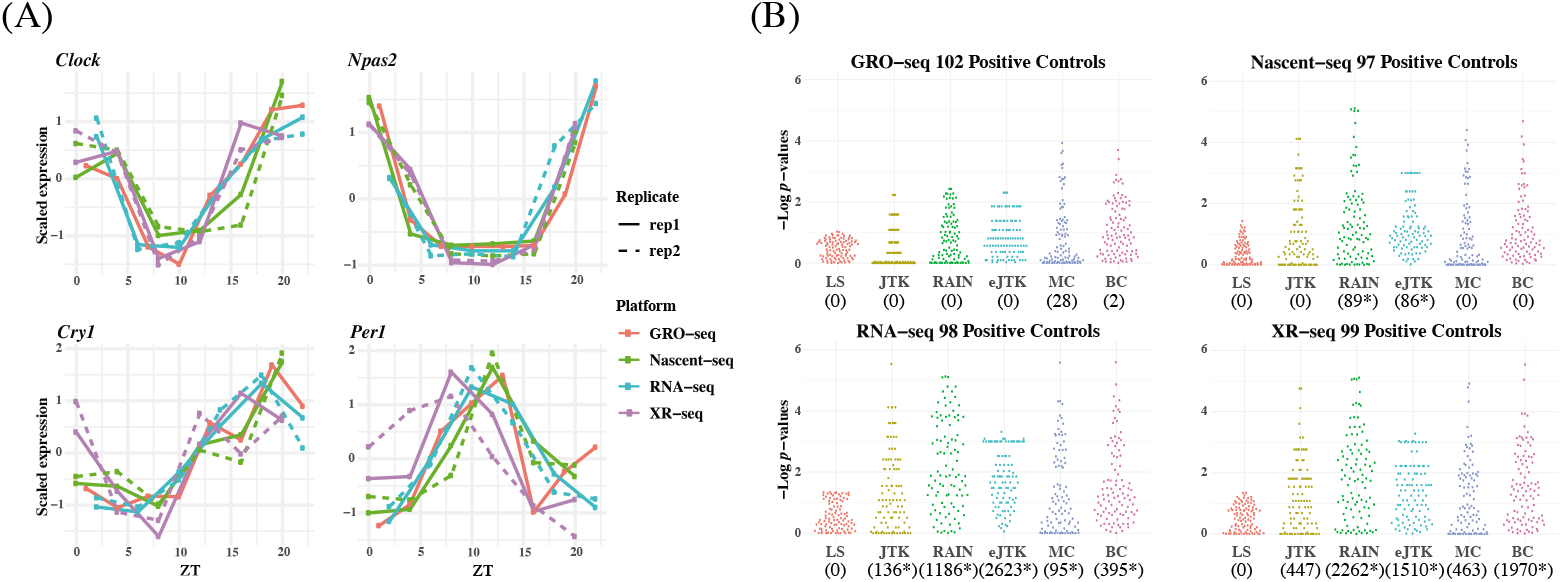
Circadian rhythm detection under light-dark experimental design by GRO-seq, Nascent-seq, RNA-seq, and XR-seq. (A) Gene-specific measurements of nascent RNA, RNA, and transcription-coupled repair of four circadian benchmark genes, *Clock, Npas2, Cry1*, and *Per1* by four different sequencing platforms. The solid and dotted lines are used for the first and second replicates respectively. (B) Beehive plots of negative log *p*-values of base 10 of circadian genes as positive controls. The number of significant genes detected by each method with an FDR threshold of 0.05 are shown in parenthesis. The asterisks denote significant GO enrichments of circadian rhythm pathway. The nominal *p*-values by JTK_CYCLE, MetaCycle, and BIO_CYCLE are the most significant, while LS and RAIN tend to be underpowered. ARS is not included in the analysis because it cannot be applied to datasets with replicates.

ARSER, despite its high reproducibility, cannot handle replicates, and previous studies have shown that data should never be concatenated [17]. Therefore, we focused on assessing the performance of the other six methods. We first examined the distribution of the nominal *p*-values of the 104 gold-standard circadian genes returned by each method, visualized as beehive plots in Figure 4B, where LS is significantly underpowered in the detection of circadian genes compared to the other methods, given any of the sequencing platforms. This result can be attributed to LS’s inability to effectively detect circadian rhythms in datasets with low sampling resolution, which is concordant with our previous results. We observe that JTK_CYCLE, RAIN, eJTK_CYCLE, MetaCycle, and BIO_CYCLE can withstand the sparse sampling and result in overall good performance.

To further assess the performance of the methods, we examined the number of significant genes identified by each method with a false discovery rate (FDR) of 0.05. Of the 9,481 mutual genes in the four datasets, LS did not identify any significant genes in any of the datasets. This result aligns with the results from the previous analysis, where we observed LS as being underpowered. JTK_CYCLE and MetaCycle detected a relatively small number of significant genes by RNA-seq and XR-seq. eJTK_CYCLE identified 2,623 significant genes by RNA-seq, and RAIN and BIO_CYCLE identified 2,262 and 1,970 significant genes by XR-seq, respectively. When comparing across different sequencing platforms, we observe that the number of detected significant genes from RNA-seq and XR-seq data is much higher than that of the GRO-seq and Nascent-seq data. This implicates a potential deficiency in detecting gene expression rhythmicity by measuring nascent transcripts.

With the identified significant genes, we further carried out a gene set enrichment analysis using the DAVID web server [25, 26] with the default options. Results from the KEGG pathway enrichment analysis are shown in Supplementary Table 2. We find that circadian rhythm is significantly enriched by various algorithms, which are marked with asterisks in Figure 4B. Specifically, we find that of the five methods that were able to identify statistically significant genes from RNA-seq data, all have enriched circadian rhythm pathway. Circadian rhythm is also enriched in the three lists of genes that were identified by eJTK_CYCLE and RAIN as well as two of the three lists of genes identified by BIO_CYCLE.

### Performance assessment using empirical proteomic dataset of dark-dark design

To assess performance of the various methods on non-transcriptomic data, we adopted a proteomic dataset of mouse livers under dark-dark experimental design from Robles et. al [27]. Refer to the Materials and Methods section for details. Since this dataset consists of replicates and missing values, only LS, JTK_CYCLE, RAIN, and MetaCycle were directly applicable. eJTK_CYCLE was not included due to its inefficiency in handling random missing values across different genes/proteins. We calculated the number of significant proteins identified by each method using an FDR threshold of 0.05 (Supplementary Figure 1A). LS identified the least number of oscillatory proteins. JTK_CYCLE and MetaCycle returned a moderate number of significant proteins. RAIN identified the largest number of oscillatory proteins, 582, exceeding that of other methods by more than 300. Heatmaps of scaled measurements of oscillatory proteins identified by at least two methods are shown in Supplementary Figure 1B, where the proteins are ordered based on their inferred phases. With the identified oscillatory proteins, we conducted a gene set enrichment analysis using the DAVID web server. While the results did not indicate that circadian rhythm was significantly enriched by any of the algorithms, KEGG metabolic pathways were significantly enriched by all algorithms but LS (Supplementary Table 3).

### Performance assessment using synthetic datasets

To provide guidelines for method selection, we evaluated the performance of the seven methods in detecting circadian rhythm by simulations with known ground truths. Examples of waveforms generated for the simulated datasets are shown in Supplementary Table 4. We generated six groups of simulated datasets to investigate how key factors affect the performance, including sampling patterns, replicates, waveforms, signal-to-noise ratios (SNRs), uneven samplings, and missing values. Supplementary Table 5 outlines the six groups of simulations and we leave the detailed setup in the Materials and Methods section. Within each simulation group, we repeated each assessment with three different sampling frequencies to determine whether increasing sampling frequency may have an effect on the aforementioned factors. The three sampling frequencies include 4 h/1 day (six time-points), 3 h/1 day (eight time-points), and 2 h/1 day (twelve time-points) and the results are shown in Figure 5A, 5B, and 5C, respectively.

**Figure 5.**
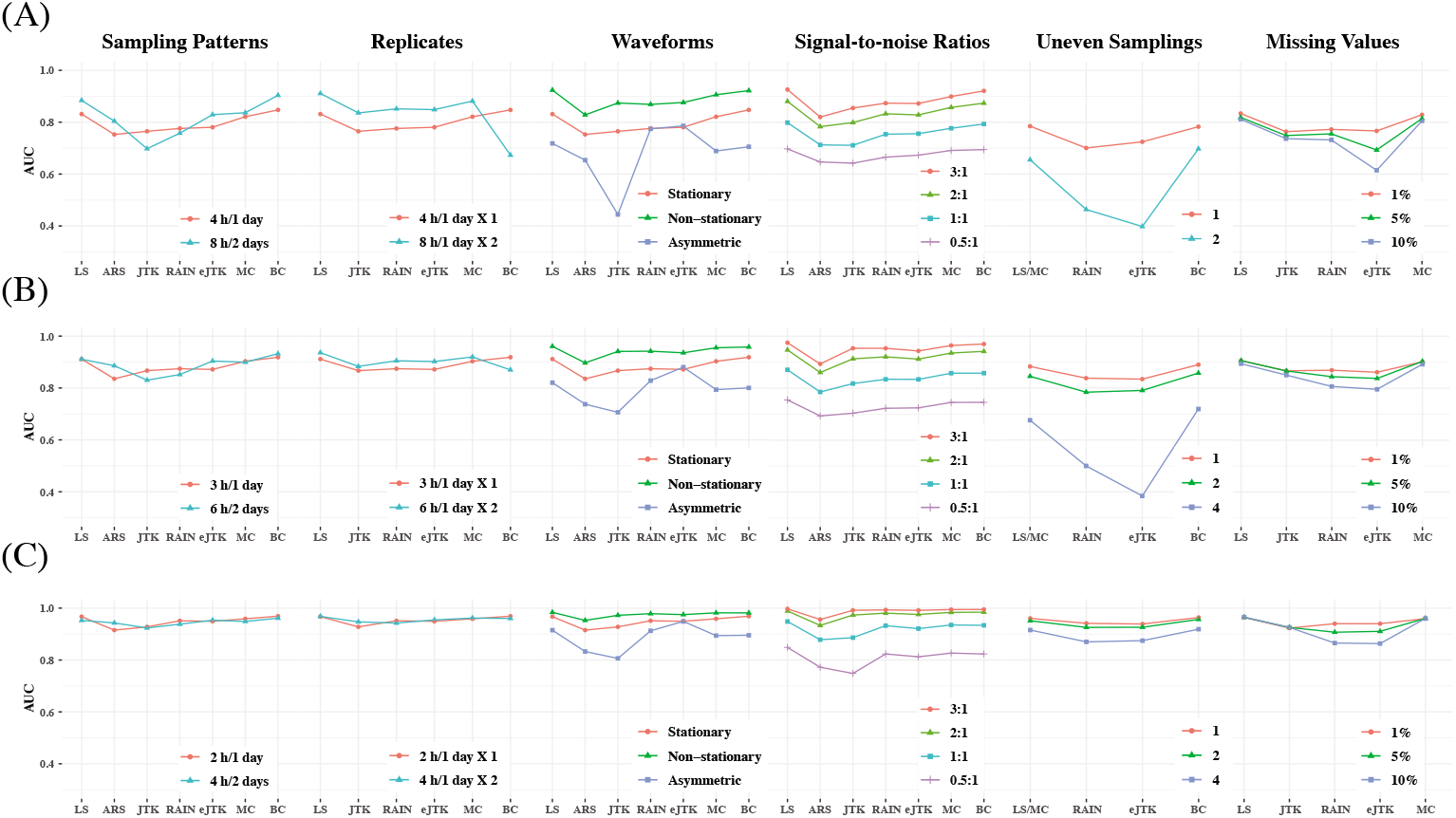
Performance assessment via simulation studies. Seven circadian rhythm detection methods are evaluated under different experimental designs to explore how sampling patterns, replicates, waveforms, signal-to-noise ratios (SNRs), uneven samplings, and missing values affect performance. Simulations under each design are carried out with different sampling frequencies: (A) 4 h/1 day, (B) 3 h/1 day, and (C) 2 h/1 day. AUC values calculated from ground truths are used as metrics.

#### Sampling patterns

To determine whether increasing the sampling frequency or lengthening the time-window is more important for each method, we first evaluated the results under the sampling pattern of 4 h/1 day versus 8 h/2 days, 3 h/1 day versus 6 h/2 days, and 2 h/1 day versus 4 h/2 days. We did not find strikingly different results within each pair of comparison, indicating that when the total number of data points are fixed, having a denser sampling density and enlarging the sampling time-window tend to have similar impact on performance. However, when we increase the number of data points, the performances of all methods are improved, which is concordant with existing studies [16, 17]. BIO_CYCLE generally outperforms the other methods, especially in datasets with lower sampling frequency and shorter time-window, while JTK_CYCLE is the most sensitive to fewer observations.

#### Replicates

To investigate the trade-off between replicates and sampling frequency, we compared the results of higher sampling frequency without replicates to those of lower sampling frequency with replicates. We first compared the dataset sampled at 4 h /1 day X1 to the dataset sampled at 8 h/1 day X2. LS, JTK_CYCLE, RAIN, eJTK_CYCLE, and MetaCycle show better performance with replicates, while BIO_CYCLE performs significantly better on densely sampled datasets without replicates. Similar results are seen when we applied the methods to the dataset at 3 h/1 day without replicates and the dataset at 6 h/1 day with replicates. As expected, further increasing the sampling resolution offsets the existing preferences that the methods have for inclusion of replicates or higher sampling density.

#### Waveforms

Supplementary Table 4 outlines the different types of periodic waveforms that we generated *in silico* in three broad categories: stationary, non-stationary, and asymmetric ones. Through our simulations, we find that all of the algorithms perform the best in detecting non-stationary waveforms. Additionally, all methods, with the exception of eJTK_CYCLE, perform better on stationary waveforms, compared to asymmetric waveforms. eJTK_CYCLE and RAIN are the top two methods for identifying asymmetric waveforms, which are expected due to their design. This is followed by LS, BIO_CYCLE, MetaCycle, and ARSER. JTK_CYCLE is the least effective in identifying asymmetric waveforms regardless of sampling frequency.

#### Signal-to-noise ratios (SNRs)

To test the effects of different noise levels on method performance, we generated various datasets with signal-to-noise ratios of 3, 2, 1, and 0.5. For all methods, our results suggest that the larger the SNRs, the higher the accuracy, as expected. LS, MetaCycle, and BIO_CYCLE are overall the most robust to noises regardless of sampling frequency, while JTK_CYCLE has the poorest performance given high noise levels.

#### Uneven samplings

To understand how well the methods deal with uneven samplings, we focus on the results of datasets with one or more uneven time-points. Our results suggest that BIO_CYCLE and LS/MetaCycle outperform the other two compatible methods. Under a sparse sampling design, RAIN and eJTK_CYCLE suffer significantly from an increasing number of uneven samplings; a dense sampling design, on the other hand, rescues the aforementioned methods.

#### Missing values

We generated datasets that contain 1%, 5%, and 10% missing data, and benchmarked the four methods that allow missing values. The performances of eJTK_CYCLE and RAIN degrade with an increasing proportion of missing values, while the performances of LS, JTK_CYCLE, and MetaCycle are comparably invariant, especially under dense sampling design. We note that eJTK_CYCLE does not handle missing values efficiently, unless the same sampling time points are missing across all genes, which reduces to uneven sampling. When there is not a shared missing pattern across different genes, the dataset needs to be split into multiple uneven sampling cases, and eJTK_CYCLE needs to be applied separately, followed by results integration. Note that BIO_CYCLE can be applied to datasets with missing values only if there are replicates and the missingness only pertains to part of the replicates. We therefore did not include it in the benchmark.

#### Computational efficiency

Last but not least, we evaluated the computational efficiency across all benchmarked methods. For dataset with low sampling resolution, the execution times among the methods are approximately the same (Supplementary Table 6). However, when analyzing data of larger sizes, RAIN requires significantly more time compared to the other methods. The running time for LS, ARSER, and BIO_CYCLE does not change much with varying sampling frequency. The running time for MetaCycle, which integrates results from LS, JTK_CYCLE, and ARSER, is calculated as the total running time of the three methods.

### Correlated multiple testing and non-uniform distribution of *p*-values under the null

To detect circadian rhythm across thousands of genes, multiple hypothesis testing corrections are needed [28]. A common FDR threshold of 0.05 is recommended by most methods and adjusted *p*-values (*q*-values) are returned by all methods except for RAIN. In the previous sections, we adopted both Bonferroni and Benjamini-Hochberg procedures for corrections. Here, we more carefully examine such procedures and point out a potential drawback resulted from both correlated multiple testing and non-uniform distributions of the nominal *p*-values under the null. We started with the observed expression measurements from the Hughes 2009 dataset and generated a “null” dataset by randomly permuting the time labels for each gene (Figure 6A). Such permutations not only deplete each gene’s rhythmic signals but also disrupts any gene-gene correlations as observed in the raw data, which are high between genes in the same pathways (Figure 6B). As such, all genes upon permutations are under the true null and additionally all gene-level testing is independent.

**Figure 6.**
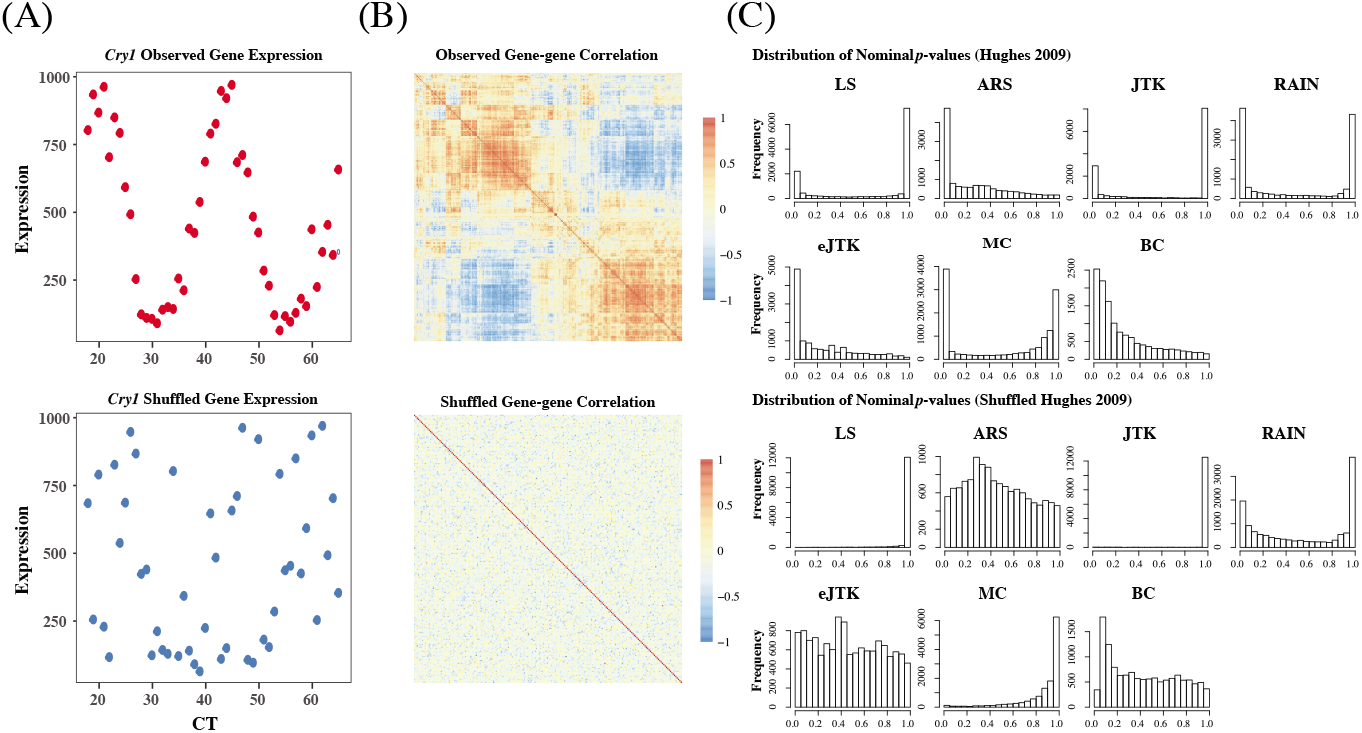
Existing methods return non-uniformly distributed *p*-values under the null, partially due to non-independent testing due to gene-gene correlations. (A) Gene expression values for the benchmark circadian gene *Cry1* before and after random permutations of the time labels. (B) Heatmaps of pairwise correlation coefficients among the top 200 highly variable genes from the Hughes 2009 dataset. The top illustrates the gene-gene correlation coefficients calculated from raw data input, and the bottom shows the gene-gene correlations after permutation. (C) The distributions of nominal *p*-values for each method when applied to the dataset before and after permutation. Gene-gene correlations, which are accounted for by eJTK_CYCLE, partially lead to the systematic deviations from the null distributions. The hypothesis testing by LS, JTK_CYCLE, RAIN, and MetaCycle are overly conservative, while ARSER’s and BIO_CYCLE’s testing procedures are biased with an overabundance of *p*-values around 0.3 and 0.1, respectively, under the null.

Figure 6C shows the distributions of nominal *p*-values for each method when applied to the dataset before and after permutation. The “U-shaped” histograms of the *p*-values for LS, JTK_CYCLE, MetaCycle, and RAIN using the original data indicate that there is dependence among the variables in the data. This violates the underlying assumption of uniformity and raises a red flag for using Bonferroni or FDR for error control [28]. A few methods have been developed for *p*-value adjustment when the tests are correlated [29–31] and such issue has been specifically pointed out by Hutchison and Dinner [32] for circadian rhythm detection.

We further applied the methods to the permuted data without gene-gene correlations. The hypothesis testing by LS, JTK_CYCLE, RAIN, and MetaCycle are still overly conservative, while the testing procedures for ARSER and BIO_CYCLE are biased with an overabundance of *p*-values around 0.3 and 0.1, respectively. eJTK_CYCLE empirically calculates the null distribution of the *p*-values via permutations and its enhanced version, booteJTK, speeds up this calculation by approximating the null distribution of the Kendall’s tau using a Gamma distribution [33]. This indeed leads to a *p*-value distribution closest to the null. However, neither eJTK_CYCLE nor booteJTK handles missing values efficiently, as explained previously. As a summary, there is still room for method development to yield *p*-values that better match the underlying assumption of a uniformly distributed *p*-values under the null.

## DISCUSSION

Here, we propose a benchmark framework to systematically evaluate the performance of seven circadian rhythm detection methods, using high-throughput omics data. The empirical datasets that we adopted in this paper were from microarray [18–20] and RNA-seq [22] to measure gene expression, Nascent-seq [22] and GRO-seq [23] to measure nascent RNA, and XR-seq [24] to measure transcription-coupled repair. While these omics data were generated from different platforms, they focus on directly or indirectly profiling transcription. It has been well studied that biological rhythm goes beyond the transcriptomic transcript-level oscillations [34]. For example, post-translational protein acetylation has been linked to circadian rhythm via mass spectrometry [35, 36]. Moreover, it has been shown that a large number of metabolites and proteins exhibit circadian oscillations [27, 37, 38]. The methods and the evaluation procedures are not limited to transcriptomic studies, but can also be applied to acetylomic, metabolomic, and proteomic experiments.

Given the assessment results from both simulations and empirical dataset anaylsis, as well as literature review of the seven methods, we have summarized the strengths and weaknesses of each method in Table 3. In general, LS, RAIN, eJTK_CYCLE, and MetaCycle are more versatile in that they can be applied to datasets with replicates, uneven samplings, or missing values. eJTK_CYCLE and BIO_CYCLE generally outperform the other methods under most situations except for handling missing values. On the other hand, JTK is sensitive to high noise levels and low sampling resolutions, and LS cannot detect any significant genes when sampling resolution is lower than 2 h/2 days with an FDR threshold of 0.05. The best detection algorithm depends on experimental designs and characteristics of the input data. Therefore, we have created two decision trees, one for low sampling resolution and the other for high sampling resolution, that outline the recommended method(s) under different scenarios (Supplementary Figure 2).

**Table 3.** Pros and cons of circadian rhythm detection methods.

| Methods | Pros | Cons |
| --- | --- | --- |
| **LS** | • Effective in handling missing values<br>• Not restricted by input data structure (i.e. can be applied to datasets with replicates, uneven samplings, or missing values) | • Rapid degradation in detectability when applied to datasets with low sampling resolution<br>• U-shaped $p$-values distribution<br>• Sensitive to outliers |
| **ARSER** | • High reproducibility | • Cannot handle replicates, uneven samplings, or missing values |
| **JTK_CYCLE** | • High precision<br>• Robust to outliers | • Incapable of detecting asymmetric waveforms<br>• U-shaped $p$-values distribution<br>• Sensitive to high level of noise<br>• High false negative rates<br>• Low reproducibility |
| **RAIN** | • High recall<br>• Effective in detecting asymmetric waveforms<br>• High reproducibility<br>• Not restricted by input data structure | • High false positive rates<br>• U-shaped $p$-values distribution<br>• Computationally intensive with increasing sampling resolution |
| **eJTK_CYCLE** | • Uniform distribution of nominal $p$-values<br>• Most effective in detecting asymmetric waveforms | • Unable to test different periods simultaneously<br>• Inefficient in handling missing values<br>• Sensitive to high level of uneven samplings |
| **MetaCycle** | • High recall<br>• Not restricted by input data structure<br>• Offset the disadvantages of one method with the other two among LS, ARSER and JTK_CYCLE<br>• Directly return calling results from three perspective methods and perform ensemble | • $P$-values generated with Fisher's integration require independence assumption |
| **BIO_CYCLE** | • Most effective in controlling for false positive rates<br>• Most robust to data with high noise, uneven samplings, and low sampling resolutions.<br>• High precision<br>• High computational efficiency with pre-trained model | • Require extensive time to train the DNN model<br>• Handle missing values only if data have replicates and the missingness only pertains to part of the replicates.<br>• Low reproducibility |

Recent advances of high-throughput technologies enable circadian rhythm detection on the genome-wide scale. As with all genomic data, the multi-time-point omics data for circadian rhythm detection bear both technical and biological variability, which can bias the analysis if not properly accounted. Data normalization and batch effect correction are crucial to remove technical biases and artifacts [39]. Cross-subject variability in rhythmic profiles, especially for human subjects, is a non-negligible source of genetic variation that needs to be adjusted [14]. This is especially important in the case-control setting where multiple subjects are involved. While we did not particularly focus on differential analysis since it is outside the scope of this paper, a few methods, including LimoRhyde [40] and DODR [12] have been made available for differential rhythmicity analysis under different conditions.

Increasingly more circadian omics data are being made available through existing studies and databases [34, 41]. We showed, from our empirical studies, that the rhythmic signals can be well recapitulated across different studies and/or different platforms (Figure 1, Figure 4A). Meta-analysis and multi-omics data integration remain an open-ended question in circadian rhythm detection [42]. In addition, transfer learning has been applied to multiple genomic research domains in genomics [43] – to borrow information and to transfer knowledge from existing data deposited in public repositories remain one of the future directions. Similarly, across different methods, an ensemble framework, as implemented by MetaCycle, can potentially boost performance. However, as we have pointed out earlier, the instability issue needs to be addressed, especially when multiple drastically distinct results are to be integrated.

To our best knowledge, all existing studies for circadian rhythm detection resort to bulk-tissue omics data, which characterize an averaged profile across different cell types in a tissue. The inherent heterogeneity can bias the analysis with reduced power and/or inflated FDR. Single-cell sequencing circumvents the averaging artifacts associated with traditional bulk population data and has seen rapid technological developments over the past few years. To assess the feasibility of single-cell circadian rhythm detection, we *in silico* generated single-cell RNA sequencing profiles by downsampling bulk RNA-seq read counts. Gold-standard circadian and noncircadian genes were used to calculate the associated AUC values (Supplementary Table 7). All methods suffer from low sequencing depth – a characteristic of the single-cell data. With the decreasing cost and the increasing popularity of single-cell omics techniques, to profile circadian rhythmicity at the cellular level and to disentangle within tissue heterogeneity with regard to biological rhythm can be of great impact.

## MATERIALS AND METHODS

### Empirical transcriptomic datasets

Three datasets under the dark-dark experimental design including Hughes 2009 [18], Hughes 2012 [19], and Zhang 2014 [20] were downloaded from GEO, and all used microarrays to profile gene expressions (Table 2A). Additionally, we obtained four datasets under the light-dark experimental design from the different sequencing platforms, including Nascent-sequencing (Nascent-seq) [22], RNA-sequencing (RNA-seq) [22], Global Run-On sequencing (GRO-seq) [23], and eXcision Repair-sequencing (XR-seq) [24] (Table 2B). Nascent-seq sequence transcribed RNAs, obtained from the nuclei without formation of the 3’ end [44]. GRO-seq measures nascent RNAs by mapping, characterizing, and evaluating transcriptionally engaged polymerase [45]. GRO-seq and Nascent-seq differ from traditional RNA-seq, in which the reads map to predominantly introns, while RNA-seq mainly assays exons [44]. XR-seq profiles DNA excision repair on the genome-wide scale with single-nucleotide resolution [46]. Here, we focus on XR-seq data from the transcribed strand only – it has been shown that the transcription-coupled repair from the transcribed strand is positively correlated with expression [47].

For quality control, we removed genes that had constant gene expression measurements in all datasets and further removed genes with more than half zero gene expression values in the light-dark datasets. In cases where multiple probes got mapped to the same RefSeq loci, we averaged the gene expression of the probes using the limma package [48], available in Bioconductor. For data normalization, robust multi-array average (RMA) [49] and genechip RMA (GC-RMA) [50] were used to normalize the array data; transcript per million (TPM) and reads per kilobase per million reads (RPKM) [51] were used to normalize the transcriptomic sequencing data. We scaled the normalized data within each gene to make them compatible for visualization only, as shown in Figure 1 and Figure 4A.

### Empirical proteomic dataset

A proteomic dataset of *Mus musculus* liver tissues from Robles et. al [27] was adopted to detect oscillatory proteins. Mouse liver samples were collected from a total of 64 mice that were released into constant darkness for one day after being entrained to a 12-12 hour light-dark schedule for 10 days. Four mice were sacrificed every 3 hours for 2 days. Then, in vivo Stable Isotope Labeling by Amino acids in Cell culture (SILAC) [52, 53] in combination with mass spectrometry was performed to profile the proteome. For each time point, equal amount of protein liver extracts from the four mice were mixed together with equal amount of protein lysates, collected in anti-phase, from the liver samples of two SILAC mice. The pooled protein extracts were measured with Orbitrap mass spectrometer. The protein abundance was calculated by taking the ratio of the signal for the mice and the signal for the heavy SILAC mix. After assessing quantification values, a total of 3,132 proteins remained for downstream circadian rhythm analysis.

### Downsampled RNA-seq dataset

We generated several downsampled RNA-seq datasets from the original RNA-seq dataset under the light-dark design to assess the robustness of the various methods to low sequencing depths. We obtained the raw sequencing data from GEO, performed read alignment to the mouse reference genome (mm10) using STAR [54], carried out quality control procedures on the aligned reads, and obtained integer-valued read counts using featureCounts [55]. We then generated downsampled RNA-seq data by multinomial sampling with index 5K, 10K, 50K, 100K, and 500K, and gene-specific probability parameters calculated from the raw data. RPKM was used to normalize the downsampled RNA-seq read counts, followed by circadian rhythm detection.

### Evaluation metrics

To evaluate the performance of the benchmarked methods, we adopted a list of 104 circadian [10] and 113 non-circadian genes [21] in mouse liver as positive and negative controls, respectively. See Supplementary Table 1 for a full list of these gold-standard genes. With these gold-standard genes, we calculated metrics including the precision and recall rates given a *p*-value or *q*-value significance threshold (Figure 2A). We further calculated the AUC values of the ROC curves, as joint measures of sensitivity and specificity (Figure 2B).

To assess the reproducibility of each method, we compared the results from the four dark-dark datasets by calculating the number of overlapping genes, as well as the Jaccard and Sorensen index as metrics for similarity (Figure 3). Venn diagrams are used to display the number of overlapping cycling genes identified across different datasets by each method. The Jaccard index measures the pairwise similarities of the significant genes detected between each pair of datasets. Let *A_i_* and *A_j_* be the set of significant genes from dataset *i* and *j*. The Jaccard similarity index is defined as

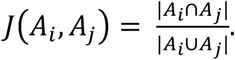

The Sorensen Index is used to characterize similarity across all datasets [56]:

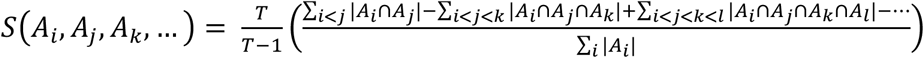

where *T* is the number of sets compared. Larger number of overlapping genes and larger Jaccard/Sorensen index values indicate higher reproducibility of the methods.

### Simulation setup

Each simulated dataset consists of 6,000 circadian and 6,000 non-circadian gene profiles. Stationary circadian profiles with a period of 24 hours are used in each simulation group, as outlined below. Note that when running the methods, we set the period range from 20 to 28 h for all methods except for eJTK_CYCLE and JTK_CYCLE, which either has a fixed period of 24 h or adjusts the period on the fly. The amplitude of the waveforms is sampled from a uniform distribution between 1 and 6; the phase shift is sampled from a uniform distribution between 0 and 24 h; and the noise term is sampled from a standard normal distribution. Flat waveforms are used to generate non-circadian profiles in all simulation groups except for testing against non-stationary waveforms where linear lines are used.

We first aimed to investigate whether higher sampling frequency or longer sampling time-window is more beneficial for each method. In this simulation group, we generated two datasets with different sampling frequencies and sampling time-windows. With six time-points, we generated one dataset at 4 h/1 day and another at 8 h/2 days; with eight time-points, we generated one dataset at 3 h/1 day and another at 6 h/2 days; with 12 time-points, we generated one dataset at 2 h/1 day and another at 4 h/2 days.

Next, we assessed whether the inclusion of replicates can offset the effect of low sampling frequency in methods’ ability of detecting oscillations. Replicates are defined as multiple measurements taken at the same time-point. Specifically, we generated two datasets consisting of the same number of observations, with or without replicates: one at 4 h/1 day X1 and the other at 8 h/1 day X2. The sampling design of the other two pairs of datasets are 3 h/1 day X1 v.s. 6 h/1 day X2, and 2 h/1 day X1 v.s. 4 h/1 day X2.

Since biological rhythms can take on various waveforms, we generated three types of waveforms via simulation: stationary, non-stationary, and asymmetric curves. Supplementary Table 4 includes models that we adopted *in silico* to generate the corresponding waveforms. Specifically, the stationary waveforms include cosine, cosine 2, and cosine peak curves; the non-stationary waveforms include cosine damp, trend exponential, and trend linear curves; the asymmetric subgroup consists of only the saw-tooth waveform. We assessed the performance of the methods in identifying each category of the circadian waveforms.

The next three groups of simulations aimed to determine which methods are more robust to different levels of signal-to-noise ratios, uneven samplings, and missing values. Specifically, we generated four datasets with SNRs of 0.5, 1, 2, and 3. Signal-to-noise ratio is defined by taking the ratio of the empirical variance of cosine function and the variance of the noise, the latter of which is fixed at one. Uneven samplings are defined as designs whose time-points are not equally spaced. To investigate the effect of uneven samplings on performance, we generated datasets with one, two, or four uneven samplings. With six time-points, datasets with four uneven samplings cannot be generated as it would only have two time-points. For missing data, we generated three levels of missing data (1%, 5%, and 10%) at three fixed, randomly selected time-points.

Lastly, we generated three datasets with sampling patterns of 1 h/2 days, 2 h/2 days, and 4 h/2 days to compute the execution times for each method. We seek to identify the differences in computational efficiency among the methods and to explore the effect of increasing sampling resolution on the execution time. Each dataset consists of a total of 6,000 genes. All execution times are reported by running on a Macbook Pro (15-inch, 2019) with 2.3 GHz 8-Core Intel Core i9 and 16 GB memory.

## DATA AND SOFTWARE AVAILABILITY

MetaCycle is an open-source R package available at https://github.com/gangwug/MetaCycle and is also used for individual analysis for LS, JTK_CYCLE, and ARSER. RAIN is a Bioconductor R package available at https://bioconductor.org/packages/rain/. eJTK_CYCLE was downloaded from https://github.com/alanlhutchison/empirical-JTK_CYCLE-with-asymmetry. BIO_CYCLE was downloaded from http://circadiomics.igb.uci.edu/BIO_CYCLE. All empirical datasets were downloaded from the NCBI Gene Expression Omnibus (https://www.ncbi.nlm.nih.gov/geo/). The accession numbers for dark-dark datasets are GSE11923, GSE30411, and GSE54652, respectively. The accession numbers for light-dark datasets are GSE59486, GSE36872, GSE36871 and GSE109938, respectively. The proteomic dataset was downloaded from the BioStudies database with accession number S-EPMC3879213.

## ACKNOWLEDGEMENTS

This work was supported by NIH Grants R35 GM118102 (to A.S.), R01 ES027255 (to A.S.), P01 CA142538 (to Y.J.), UL1 TR002489 (to Y.J.), and a pilot award from the UNC Computational Medicine Program (to Y.J.). We thank the Sancar Lab members and Dr. John Hogenesch for helpful discussions and feedback.

## SUPPLEMENTARY FIGURE & TABLE LEGENDS

**Supplementary Table 1. Circadian and non-circadian genes in *Mus muculus* liver as gold standard**. The 104 circadian gene list is extracted from Supplementary Table 4 in Wu et al. Wu G, Zhu J, Yu J, Zhou L, Huang JZ and Zhang Z [10] and the 113 non-circadian gene list is obtained from Supplementary Table 2 in Wu et al. Wu G, Zhu J, He F, Wang W, Hu S and Yu J [21].

**Supplementary Table 2. Pathway enrichment analysis of significantly cyclic genes from the light-dark datasets**. Functional annotations (KEGG pathway mapping) of the significant genes (*q*-values ≤ 0.05) are carried out using the the DAVID Bioinformatics Resources (https://david.ncifcrf.gov/). The list only contains significantly enriched pathways with a 0.05 cutoff of the *p*-values adjusted by Benjamini Hochberg.

**Supplementary Table 3. Pathway enrichment analysis of significantly cyclic proteins**. Functional annotations (KEGG pathway mapping) of the significant proteins (*q*-values ≤ 0.05) are carried out using the the DAVID Bioinformatics Resources (https://david.ncifcrf.gov/). The list only contains significantly enriched pathways with a 0.05 cutoff of the *p*-values adjusted by Benjamini Hochberg. KEGG metabolic pathways were enriched by all three methods.

**Supplementary Table 4. *In silico* generated periodic v.s. non-periodic gene profiles**. Three types of periodic waveforms are included: stationary, non-stationary, and asymmetric. The stationary and non-stationary subgroups consist of three forms of cosine curves. The asymmetric subgroup consists of a saw-tooth waveform. Flat or linear lines are adopted to generate non-periodic waveforms. The waveforms shown are constructed without noise. ‘Amp’, ‘pha’, and ‘per’ represent amplitude, phase and period, respectively.

**Supplementary Table 5. Details of simulation setup and parameters used to *in silico* generate periodic and non-periodic profiles**. Each simulation run consists of 6,000 periodic and 6,000 non-periodic gene profiles. All simulated waveforms have a period length of 24, a phase shift that is uniformly distributed between 0 and 24, and a noise term with standard normal distribution. The amplitude is uniformly distributed between 1 and 6 for all groups except when testing for different signal-to-noise ratios (SNRs), which we define as the ratios of the empirical variances of the cosine function and the variances of the noise. Non-periodic profiles are sampled from a flat/linear function. “X 1” indicates no replicate and “X 2” indicates two replicates.

**Supplementary Table 6. Evaluation of computational efficiency with different sampling rates**. Each method is run on a dataset with a total of 6,000 genes. All programs are run on a Macbook Pro (15-inch, 2019) with 2.3 GHz 8-Core Intel Core i9 and 16 GB memory. Running time for MetaCycle is the sum of the runing time for LS, ARSER, and JTK_CYCLE. Running time for BIO_CYCLE does not include the time used to fit the deep neural network.

**Supplementary Table 7. Performance assessment of downsampled RNA-seq data**. AUC values of downsampled RNA-seq datasets with varying sequencing depths were calculated. Existing methods suffer from low sequencing depths. The performance of RAIN exceeds that of all other methods in all sequencing depths with an exception at 5K, due to its large number of significant genes detected in general. BIO_CYCLE consistently ranks the lowest at all but the highest sequencing depth. The performances of LS, JTK_CYCLE, eJTK_CYCLE, and MetaCycle are comparable.

**Supplementary Figure 1. Circadian rhythm detection of *Mus musculus* liver protemoic dataset**. (A) Bar plot of the number of significant proteins detected by each method using an FDR threshold of 0.05. Only methods that are able to handle both replicates and missing values were applied and evaluated. (B) Heatmap of scaled measurements of oscillatory proteins identified by at least two methods. Proteins (rows) are ordered based on their inferred phases.

**Supplementary Figure 2. Decision tree as user guidance on method selection**. The decision tree has decision rules for sampling resolutions, uneven samplings, replicates, and missing values.

