## Supplement for "Genome-wide circadian rhythm detection methods: systematic evaluations and practical guidelines"

| Circadian Gene Symbol | | | |
|---|---|---|---|
| Acacb | Fads2 | Tgfbr1 | Fkbp5 |
| Acadm | Fasn | Alas1 | Usp2 |
| Acly | Fgf15 | Igfbp1 | Amd1 |
| Parp1 | Insig1 | Raet1c | Gck |
| Cyp2d9 | Insig2 | Rora | Hmgcr |
| Cyp7b1 | Ldlr | Sult1e1 | Lpin1 |
| Arntl | Lipg | Fas | Saa4 |
| Clock | Lpl | Rxra | Wee1 |
| Cry2 | Nr0b2 | Cirbp | Nampt |
| Npas2 | Nr1h2 | Fus | Aco2 |
| Per1 | Nr1h3 | Hsp90aa1 | Akr7a5 |
| Per2 | Pctp | Hsph1 | Aldh2 |
| Per3 | Scap | Ccrn4l | Bhmt |
| Acot1 | Srebf1 | Cpt1a | Cps1 |
| Acot2 | Srebf2 | Pklr | Eef1d |
| Acot4 | Apoc3 | Slc2a2 | Sord |
| Cyp4a10 | Avpr1a | Slc37a4 | Bhlhe41 |
| Cyp4a14 | Fkbp4 | Ucp2 | Nr1d1 |
| Ak3 | Hspd1 | Aacs | Nr1d2 |
| Rgs16 | Tubb5 | Acot3 | Rorc |
| Ccnd1 | Dbp | Apoa1 | G6pc |
| Ddc | Hlf | Cyp7a1 | Pck1 |
| Hist1h1c | Nfil3 | Elovl3 | Ppara |
| Rell1 | Tef | Elovl5 | Ppard |
| Sbk1 | Bhlhe40 | Elovl6 | Pparg |
| Smarcd1 | Por | Fabp5 | Id2 |

| Non−circadian Gene Symbol | | | |
|---|---|---|---|
| Eif2a | Zkscan14 | Trdmt1 | Zfx |
| Utp6 | Zfp1 | Tsfm | Snx12 |
| Rpl19 | Slc26a11 | Ercc6l2 | Zfp524 |
| Rplp0 | Nkap | Smg8 | Apbb3 |
| Tbp | Mobkl2c | Commd5 | Ppie |
| Eef1a1 | Eya3 | Ing5 | Cops7b |
| Hmbs | Ddx27 | Fam119a | Glmn |
| Tbcc | Asb6 | Fam175a | Zfp592 |
| Actb | Zfp451 | Nvl | Nub1 |
| Gapdh | Cln3 | Rexo1 | Zfp426 |
| Ppib | Gpn2 | Rngtt | Sertad3 |
| Polr3f | Mtf1 | Snapc3 | Zfp414 |
| Med6 | Rab23 | Cdkal1 | Axin1 |
| Tyw1 | Lysmd1 | Rrp9 | Casz1 |
| Vars2 | Asb3 | Utp15 | Bcl2l1 |
| Ell | Zfp397 | Atpbd3 | Telo2 |
| Gtf2h3 | Fam123b | Nsun5 | Terf2ip |
| Polr1a | Zkscan6 | Chd1 | Rabl4 |
| Ttf1 | Adck2 | Kdm4a | Dgcr8 |
| Mrps5 | Usp36 | Men1 | Cep110 |
| Med23 | Trib1 | Kdm5d | Abcd4 |
| Polr3h | Tfpt | Ankra2 | Rad50 |
| Patl1 | Stam2 | Bcor | Sesn2 |
| Med24 | E2f3 | Rbm28 | Gnptab |
| Tsr3 | Zfp511 | Kat2a | Gtf2ird2 |
| Rprd2 | Slc10a7 | Gtpbp5 | Zscan12 |
| Tbc1d5 | Vps33b | Mynn | Ino80b |
| Ttc15 | Ppm1f | Sumf2 | Smurf1 |
| Mga | | | |

Supplementary Table 1

| Method | Term | Count | Percent | $P$−Value | Benjamini |
|---|---|---|---|---|---|
| GRO−seq MC | mmu00260:Glycine, serine and threonine metabolism | 3 | 11.5384615 | 1.408300e−03 | 4.812851e−02 |
| Nascent−seq RAIN | mmu04710:Circadian rhythm | 4 | 4.4943820 | 4.546934e−04 | 4.359144e−02 |
| Nascent−seq eJTK | mmu04710:Circadian rhythm | 5 | 5.8823529 | 1.448933e−05 | 1.259787e−03 |
| RNA−seq JTK | mmu04710:Circadian rhythm | 5 | 3.7037037 | 1.738213e−04 | 2.064430e−02 |
| RNA−seq RAIN | mmu01100:Metabolic pathways | 169 | 14.2978003 | 8.003523e−16 | 2.176037e−13 |
| | mmu00982:Drug metabolism – cytochrome P450 | 25 | 2.1150592 | 1.644394e−11 | 2.302154e−09 |
| | mmu00980:Metabolism of xenobiotics by cytochrome P450 | 24 | 2.0304569 | 5.798815e−11 | 5.412229e−09 |
| | mmu00040:Pentose and glucuronate interconversions | 16 | 1.3536379 | 2.901869e−09 | 2.031308e−07 |
| | mmu05204:Chemical carcinogenesis | 26 | 2.1996616 | 6.999948e−09 | 3.919970e−07 |
| | mmu00860:Porphyrin and chlorophyll metabolism | 16 | 1.3536379 | 1.012513e−07 | 4.725049e−06 |
| | mmu00053:Ascorbate and aldarate metabolism | 12 | 1.0152284 | 1.519743e−06 | 6.078791e−05 |
| | mmu00983:Drug metabolism – other enzymes | 16 | 1.3536379 | 2.527470e−06 | 8.845766e−05 |
| | mmu04710:Circadian rhythm | 11 | 0.9306261 | 5.067967e−05 | 1.575498e−03 |
| | mmu00830:Retinol metabolism | 19 | 1.6074450 | 7.641930e−05 | 2.137534e−03 |
| | mmu01130:Biosynthesis of antibiotics | 31 | 2.6226734 | 5.198836e−04 | 1.314962e−02 |
| | mmu00480:Glutathione metabolism | 13 | 1.0998308 | 5.774587e−04 | 1.338751e−02 |
| | mmu04141:Protein processing in endoplasmic reticulum | 26 | 2.1996616 | 6.167556e−04 | 1.320017e−02 |
| | mmu00130:Ubiquinone and other terpenoid−quinone biosynthesis | 6 | 0.5076142 | 7.081090e−04 | 1.406731e−02 |
| | mmu04146:Peroxisome | 16 | 1.3536379 | 1.041198e−03 | 1.925798e−02 |
| | mmu00140:Steroid hormone biosynthesis | 16 | 1.3536379 | 1.708722e−03 | 2.948479e−02 |
| | mmu00270:Cysteine and methionine metabolism | 10 | 0.8460237 | 2.213890e−03 | 3.584623e−02 |
| RNA−seq eJTK | mmu01100:Metabolic pathways | 321 | 12.2753346 | 5.097538e−24 | 1.478286e−21 |
| | mmu00982:Drug metabolism – cytochrome P450 | 35 | 1.3384321 | 7.422532e−12 | 1.076264e−09 |
| | mmu00980:Metabolism of xenobiotics by cytochrome P450 | 34 | 1.3001912 | 1.466155e−11 | 1.417289e−09 |
| | mmu01130:Biosynthesis of antibiotics | 70 | 2.6768642 | 3.479491e−10 | 2.522630e−08 |
| | mmu00280:Valine, leucine and isoleucine degradation | 27 | 1.0325048 | 2.296309e−08 | 1.331858e−06 |
| | mmu05204:Chemical carcinogenesis | 37 | 1.4149140 | 2.547489e−08 | 1.231285e−06 |
| | mmu00071:Fatty acid degradation | 25 | 0.9560229 | 3.356184e−08 | 1.390418e−06 |
| | mmu01200:Carbon metabolism | 40 | 1.5296367 | 7.604773e−07 | 2.756693e−05 |
| | mmu00053:Ascorbate and aldarate metabolism | 16 | 0.6118547 | 1.688110e−06 | 5.439322e−05 |
| | mmu04146:Peroxisome | 31 | 1.1854685 | 2.657397e−06 | 7.706165e−05 |
| | mmu00040:Pentose and glucuronate interconversions | 17 | 0.6500956 | 7.960225e−06 | 2.098393e−04 |
| | mmu00860:Porphyrin and chlorophyll metabolism | 19 | 0.7265774 | 1.246414e−05 | 3.011731e−04 |
| | mmu00830:Retinol metabolism | 31 | 1.1854685 | 1.338463e−05 | 2.985377e−04 |
| | mmu03050:Proteasome | 20 | 0.7648184 | 1.425353e−05 | 2.952102e−04 |
| | mmu05012:Parkinson's disease | 44 | 1.6826004 | 1.880681e−05 | 3.635357e−04 |
| | mmu04932:Non−alcoholic fatty liver disease (NAFLD) | 45 | 1.7208413 | 3.346485e−05 | 6.063766e−04 |
| | mmu05010:Alzheimer's disease | 49 | 1.8738050 | 3.797554e−05 | 6.476206e−04 |
| | mmu04710:Circadian rhythm | 15 | 0.5736138 | 7.954074e−05 | 1.280720e−03 |
| | mmu01230:Biosynthesis of amino acids | 26 | 0.9942639 | 1.084256e−04 | 1.653637e−03 |
| | mmu04610:Complement and coagulation cascades | 26 | 0.9942639 | 1.084256e−04 | 1.653637e−03 |
| | mmu05230:Central carbon metabolism in cancer | 23 | 0.8795411 | 1.339332e−04 | 1.940277e−03 |
| | mmu00270:Cysteine and methionine metabolism | 17 | 0.6500956 | 1.469660e−04 | 2.027621e−03 |
| | mmu04919:Thyroid hormone signaling pathway | 34 | 1.3001912 | 1.606747e−04 | 2.115913e−03 |
| | mmu00190:Oxidative phosphorylation | 39 | 1.4913958 | 1.987891e−04 | 2.503581e−03 |
| | mmu01040:Biosynthesis of unsaturated fatty acids | 13 | 0.4971319 | 3.111264e−04 | 3.752969e−03 |
| | mmu05016:Huntington's disease | 50 | 1.9120459 | 3.643095e−04 | 4.217841e−03 |
| | mmu00983:Drug metabolism – other enzymes | 19 | 0.7265774 | 3.704177e−04 | 4.123821e−03 |
| | mmu01212:Fatty acid metabolism | 19 | 0.7265774 | 3.704177e−04 | 4.123821e−03 |
| | mmu00130:Ubiquinone and other terpenoid−quinone biosynthesis | 8 | 0.3059273 | 3.795185e−04 | 4.068784e−03 |
| | mmu00062:Fatty acid elongation | 12 | 0.4588910 | 9.205811e−04 | 9.493629e−03 |
| | mmu00480:Glutathione metabolism | 19 | 0.7265774 | 1.038016e−03 | 1.033181e−02 |
| | mmu00140:Steroid hormone biosynthesis | 26 | 0.9942639 | 1.085668e−03 | 1.044556e−02 |
| | mmu04914:Progesterone−mediated oocyte maturation | 26 | 0.9942639 | 1.085668e−03 | 1.044556e−02 |
| | mmu00380:Tryptophan metabolism | 17 | 0.6500956 | 1.212200e−03 | 1.128268e−02 |
| | mmu00650:Butanoate metabolism | 12 | 0.4588910 | 1.339581e−03 | 1.207460e−02 |
| | mmu04141:Protein processing in endoplasmic reticulum | 42 | 1.6061185 | 1.425778e−03 | 1.246023e−02 |
| | mmu00010:Glycolysis / Gluconeogenesis | 21 | 0.8030593 | 1.643029e−03 | 1.392769e−02 |
| | mmu00260:Glycine, serine and threonine metabolism | 15 | 0.5736138 | 1.788668e−03 | 1.472418e−02 |
| | mmu00410:beta−Alanine metabolism | 13 | 0.4971319 | 2.595550e−03 | 2.071815e−02 |
| | mmu00310:Lysine degradation | 17 | 0.6500956 | 3.923485e−03 | 3.034227e−02 |
| | mmu04976:Bile secretion | 21 | 0.8030593 | 4.242325e−03 | 3.192383e−02 |
| | mmu01210:2−Oxocarboxylic acid metabolism | 9 | 0.3441683 | 4.810340e−03 | 3.522031e−02 |
| | mmu00640:Propanoate metabolism | 11 | 0.4206501 | 5.011554e−03 | 3.576970e−02 |
| | mmu00340:Histidine metabolism | 10 | 0.3824092 | 5.039932e−03 | 3.510736e−02 |
| | mmu04915:Estrogen signaling pathway | 26 | 0.9942639 | 6.271672e−03 | 4.251073e−02 |
| | mmu00020:Citrate cycle (TCA cycle) | 12 | 0.4588910 | 6.343808e−03 | 4.201208e−02 |
| | mmu02010:ABC transporters | 15 | 0.5736138 | 7.532058e−03 | 4.860982e−02 |
| RNA−seq MC | mmu04710:Circadian rhythm | 5 | 5.3191489 | 6.382092e−05 | 6.235130e−03 |
| RNA−seq BC | mmu04710:Circadian rhythm | 9 | 2.2842640 | 6.040930e−07 | 1.413478e−04 |
| | mmu01100:Metabolic pathways | 52 | 13.1979695 | 2.048620e−04 | 2.368628e−02 |

Supplements Table 2

| Category | Term | Count | Percent | $P$−Value | Benjamini |
|---|---|---|---|---|---|
| XR−seq RAIN | mmu01100:Metabolic pathways | 230 | 10.1769912 | 5.399408e−09 | 1.538830e−06 |
| | mmu04146:Peroxisome | 29 | 1.2831858 | 7.423091e−07 | 1.057735e−04 |
| | mmu04710:Circadian rhythm | 15 | 0.6637168 | 1.013260e−05 | 9.621386e−04 |
| | mmu04141:Protein processing in endoplasmic reticulum | 43 | 1.9026549 | 1.257843e−05 | 8.958171e−04 |
| | mmu03040:Spliceosome | 36 | 1.5929204 | 2.050925e−05 | 1.168356e−03 |
| | mmu03050:Proteasome | 18 | 0.7964602 | 2.103473e−05 | 9.986614e−04 |
| | mmu04152:AMPK signaling pathway | 35 | 1.5486726 | 2.170888e−05 | 8.834807e−04 |
| | mmu05169:Epstein−Barr virus infection | 35 | 1.5486726 | 8.306604e−05 | 2.954976e−03 |
| | mmu03013:RNA transport | 41 | 1.8141593 | 8.816172e−05 | 2.788017e−03 |
| | mmu00270:Cysteine and methionine metabolism | 15 | 0.6637168 | 2.880869e−04 | 8.178035e−03 |
| | mmu03010:Ribosome | 35 | 1.5486726 | 3.128193e−04 | 8.073366e−03 |
| | mmu04068:FoxO signaling pathway | 33 | 1.4601770 | 3.284607e−04 | 7.771865e−03 |
| | mmu04932:Non−alcoholic fatty liver disease (NAFLD) | 37 | 1.6371681 | 3.354265e−04 | 7.327834e−03 |
| | mmu01130:Biosynthesis of antibiotics | 46 | 2.0353982 | 5.152173e−04 | 1.043622e−02 |
| | mmu04920:Adipocytokine signaling pathway | 21 | 0.9292035 | 5.603045e−04 | 1.059227e−02 |
| | mmu04120:Ubiquitin mediated proteolysis | 32 | 1.4159292 | 1.972447e−03 | 3.455766e−02 |
| XR−seq eJTK | mmu01100:Metabolic pathways | 153 | 10.1526211 | 2.637646e−06 | 7.356334e−04 |
| | mmu04710:Circadian rhythm | 12 | 0.7962840 | 2.807031e−05 | 3.908206e−03 |
| | mmu04152:AMPK signaling pathway | 25 | 1.6589250 | 1.780315e−04 | 1.642207e−02 |
| | mmu03040:Spliceosome | 25 | 1.6589250 | 3.253113e−04 | 2.243858e−02 |
| | mmu03013:RNA transport | 29 | 1.9243530 | 5.151755e−04 | 2.834473e−02 |
| | mmu04120:Ubiquitin mediated proteolysis | 25 | 1.6589250 | 8.709083e−04 | 3.970512e−02 |
| | mmu04068:FoxO signaling pathway | 24 | 1.5925680 | 8.906319e−04 | 3.489064e−02 |
| | mmu03050:Proteasome | 12 | 0.7962840 | 1.113533e−03 | 3.811088e−02 |
| | mmu01130:Biosynthesis of antibiotics | 33 | 2.1897810 | 1.190583e−03 | 3.625647e−02 |
| XR−seq BC | mmu01100:Metabolic pathways | 200 | 10.1781170 | 5.203804e−08 | 1.488277e−05 |
| | mmu03013:RNA transport | 42 | 2.1374046 | 1.096642e−06 | 1.568075e−04 |
| | mmu03050:Proteasome | 17 | 0.8651399 | 1.435104e−05 | 1.367207e−03 |
| | mmu04152:AMPK signaling pathway | 32 | 1.6284987 | 2.042087e−05 | 1.459042e−03 |
| | mmu01212:Fatty acid metabolism | 18 | 0.9160305 | 2.073602e−05 | 1.185410e−03 |
| | mmu04146:Peroxisome | 24 | 1.2213740 | 2.445182e−05 | 1.164872e−03 |
| | mmu04141:Protein processing in endoplasmic reticulum | 38 | 1.9338422 | 3.290665e−05 | 1.343590e−03 |
| | mmu01130:Biosynthesis of antibiotics | 45 | 2.2900763 | 3.714389e−05 | 1.327037e−03 |
| | mmu04710:Circadian rhythm | 13 | 0.6615776 | 6.531774e−05 | 2.073567e−03 |
| | mmu04920:Adipocytokine signaling pathway | 20 | 1.0178117 | 2.475823e−04 | 7.056713e−03 |
| | mmu03040:Spliceosome | 29 | 1.4758270 | 6.326819e−04 | 1.632029e−02 |
| | mmu00071:Fatty acid degradation | 15 | 0.7633588 | 6.711375e−04 | 1.587348e−02 |
| | mmu03015:mRNA surveillance pathway | 23 | 1.1704835 | 7.180379e−04 | 1.567830e−02 |
| | mmu04530:Tight junction | 22 | 1.1195929 | 7.323059e−04 | 1.485402e−02 |
| | mmu04932:Non−alcoholic fatty liver disease (NAFLD) | 32 | 1.6284987 | 1.041217e−03 | 1.966692e−02 |
| | mmu04140:Regulation of autophagy | 10 | 0.5089059 | 1.391158e−03 | 2.457719e−02 |
| | mmu04120:Ubiquitin mediated proteolysis | 29 | 1.4758270 | 1.823600e−03 | 3.024072e−02 |
| | mmu05100:Bacterial invasion of epithelial cells | 19 | 0.9669211 | 1.954260e−03 | 3.060334e−02 |
| | mmu04068:FoxO signaling pathway | 27 | 1.3740458 | 3.291902e−03 | 4.842192e−02 |

Supplements Table 2 cont.

| Method | Term | Count | Percent | $P$-Value | Benjamini |
|---|---|---|---|---|---|
| JTK | mmu04610:Complement and coagulation cascades | 8 | 7.4074074 | 6.721172e–06 | 7.256256e–04 |
| | mmu01100:Metabolic pathways | 25 | 23.1481481 | 4.274151e–04 | 2.282092e–02 |
| RAIN | mmu01100:Metabolic pathways | 173 | 32.6415094 | 1.199819e–40 | 2.567613e–38 |
| | mmu01130:Biosynthesis of antibiotics | 56 | 10.5660377 | 3.862699e–25 | 4.133088e–23 |
| | mmu01200:Carbon metabolism | 36 | 6.7924528 | 9.865999e–19 | 7.037746e–17 |
| | mmu00071:Fatty acid degradation | 24 | 4.5283019 | 1.203313e–17 | 6.437722e–16 |
| | mmu01212:Fatty acid metabolism | 21 | 3.9622642 | 1.230112e–13 | 5.264900e–12 |
| | mmu00620:Pyruvate metabolism | 18 | 3.3962264 | 1.108719e–12 | 3.954248e–11 |
| | mmu00280:Valine, leucine and isoleucine degradation | 19 | 3.5849057 | 7.498068e–11 | 2.292267e–09 |
| | mmu00020:Citrate cycle (TCA cycle) | 15 | 2.8301887 | 1.088804e–10 | 2.912552e–09 |
| | mmu04141:Protein processing in endoplasmic reticulum | 31 | 5.8490566 | 8.613214e–10 | 2.048031e–08 |
| | mmu00190:Oxidative phosphorylation | 28 | 5.2830189 | 8.951829e–10 | 1.915691e–08 |
| | mmu00830:Retinol metabolism | 22 | 4.1509434 | 1.760418e–09 | 3.424813e–08 |
| | mmu00640:Propanoate metabolism | 13 | 2.4528302 | 1.948424e–09 | 3.474690e–08 |
| | mmu04146:Peroxisome | 21 | 3.9622642 | 2.917345e–09 | 4.802398e–08 |
| | mmu01230:Biosynthesis of amino acids | 20 | 3.7735849 | 3.764751e–09 | 5.754691e–08 |
| | mmu03320:PPAR signaling pathway | 20 | 3.7735849 | 9.469415e–09 | 1.350970e–07 |
| | mmu05012:Parkinson's disease | 26 | 4.9056604 | 8.303846e–08 | 1.110639e–06 |
| | mmu00380:Tryptophan metabolism | 14 | 2.6415094 | 3.196986e–07 | 4.024434e–06 |
| | mmu00630:Glyoxylate and dicarboxylate metabolism | 11 | 2.0754717 | 7.851021e–07 | 9.333951e–06 |
| | mmu00140:Steroid hormone biosynthesis | 18 | 3.3962264 | 1.153208e–06 | 1.298869e–05 |
| | mmu05010:Alzheimer's disease | 26 | 4.9056604 | 2.373644e–06 | 2.539770e–05 |
| | mmu00062:Fatty acid elongation | 10 | 1.8867925 | 2.743240e–06 | 2.795457e–05 |
| | mmu00010:Glycolysis / Gluconeogenesis | 15 | 2.8301887 | 3.714133e–06 | 3.612780e–05 |
| | mmu05016:Huntington's disease | 27 | 5.0943396 | 5.915296e–06 | 5.503662e–05 |
| | mmu05204:Chemical carcinogenesis | 17 | 3.2075472 | 1.158093e–05 | 1.032586e–04 |
| | mmu01210:2−Oxocarboxylic acid metabolism | 8 | 1.5094340 | 2.302407e–05 | 1.970689e–04 |
| | mmu01040:Biosynthesis of unsaturated fatty acids | 9 | 1.6981132 | 3.645856e–05 | 3.000424e–04 |
| | mmu00650:Butanoate metabolism | 9 | 1.6981132 | 3.645856e–05 | 3.000424e–04 |
| | mmu04932:Non−alcoholic fatty liver disease (NAFLD) | 21 | 3.9622642 | 1.093741e–04 | 8.665630e–04 |
| | mmu00220:Arginine biosynthesis | 7 | 1.3207547 | 2.420391e–04 | 1.848384e–03 |
| | mmu03050:Proteasome | 10 | 1.8867925 | 3.320438e–04 | 2.447661e–03 |
| | mmu00982:Drug metabolism – cytochrome P450 | 12 | 2.2641509 | 3.996930e–04 | 2.847651e–03 |
| | mmu00270:Cysteine and methionine metabolism | 9 | 1.6981132 | 7.105099e–04 | 4.894536e–03 |
| | mmu00310:Lysine degradation | 10 | 1.8867925 | 1.009361e–03 | 6.730756e–03 |
| | mmu00410:beta−Alanine metabolism | 8 | 1.5094340 | 1.058989e–03 | 6.847468e–03 |
| | mmu00980:Metabolism of xenobiotics by cytochrome P450 | 11 | 2.0754717 | 1.224984e–03 | 7.685233e–03 |
| | mmu00072:Synthesis and degradation of ketone bodies | 5 | 0.9433962 | 1.570289e–03 | 9.562728e–03 |
| | mmu00053:Ascorbate and aldarate metabolism | 7 | 1.3207547 | 1.875310e–03 | 1.109612e–02 |
| | mmu00330:Arginine and proline metabolism | 9 | 1.6981132 | 2.805579e–03 | 1.611835e–02 |
| | mmu00591:Linoleic acid metabolism | 9 | 1.6981132 | 3.196868e–03 | 1.787065e–02 |
| | mmu00260:Glycine, serine and threonine metabolism | 8 | 1.5094340 | 3.412465e–03 | 1.858201e–02 |
| | mmu04964:Proximal tubule bicarbonate reclamation | 6 | 1.1320755 | 4.066064e–03 | 2.156193e–02 |
| | mmu00061:Fatty acid biosynthesis | 5 | 0.9433962 | 4.223097e–03 | 2.184700e–02 |
| | mmu00360:Phenylalanine metabolism | 6 | 1.1320755 | 4.984794e–03 | 2.514080e–02 |
| | mmu00400:Phenylalanine, tyrosine and tryptophan biosynthesis | 4 | 0.7547170 | 5.857129e–03 | 2.881192e–02 |
| | mmu00120:Primary bile acid biosynthesis | 5 | 0.9433962 | 7.088942e–03 | 3.400906e–02 |
| | mmu04612:Antigen processing and presentation | 11 | 2.0754717 | 7.757074e–03 | 3.635568e–02 |
| | mmu04142:Lysosome | 14 | 2.6415094 | 8.139112e–03 | 3.730583e–02 |
| | mmu00250:Alanine, aspartate and glutamate metabolism | 7 | 1.3207547 | 9.639924e–03 | 4.314685e–02 |
| | mmu04975:Fat digestion and absorption | 7 | 1.3207547 | 1.097517e–02 | 4.801067e–02 |
| MC | mmu04610:Complement and coagulation cascades | 9 | 6.8702290 | 1.950814e–06 | 2.360209e–04 |
| | mmu01100:Metabolic pathways | 31 | 23.6641221 | 3.112798e–05 | 1.881500e–03 |
| | mmu00140:Steroid hormone biosynthesis | 7 | 5.3435115 | 4.272723e–04 | 1.708929e–02 |

Supplements Table 3

| Types | Waveforms | Equations | Examples |
|---|---|---|---|
| | | amp = A pha = $\phi$ per = T | |
| Stationary (Periodic) | cosine | $A\cos\left(\frac{2\pi(t-\phi)}{T}\right)$ | 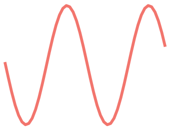 |
| | cosine 2 | $\frac{A}{1.39}\cos\left(\frac{2\pi(t-\phi_1)}{T}\right)+0.5\cos\left(\frac{2\pi(t-\phi_2)}{T_2}\right)$ where $T_2=\frac{1}{3}T$, $\phi_1=\phi+\frac{0.215T}{\pi}$, $\phi_2=(\phi_1+0.25T_2)$ mod $T$ | 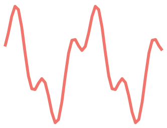 |
| | cosine peak | $A\left(-1+2\left\lvert\cos\left(\frac{\pi(t-\phi)}{T}\right)\right\rvert^{10}\right)$ | 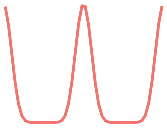 |
| Non-stationary (Periodic) | cosine damp | $A\cos\left(\frac{2\pi(t-\phi)}{T}\right)e^{-0.01t}$ | 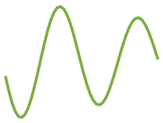 |
| | trend exp | $5e^{-0.01t}+A\cos\left(\frac{2\pi(t-\phi)}{T}\right)e^{-0.01t}$ | 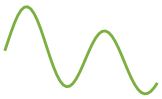 |
| | trend linear | $A\cos\left(\frac{2\pi(t-\phi)}{T}\right)+st$ where $s\sim U(-0.05,0)$ | 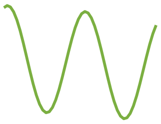 |
| Asymmetric (Periodic) | saw-tooth | $\frac{-2A}{\pi}\arctan\left(\frac{1}{\tan\frac{\pi(t-\phi)}{T}}\right)$ | 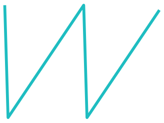 |
| Non-periodic | linear | $st$ where $s\sim U(-0.05,0)$ | 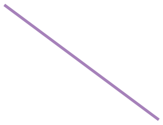 |
|  | flat | 0 |  |

Supplementary Table 4

| Simulation group | Description | Period length | Amplitude | Phase shift | Waveforms | Noise levels | Sampling patterns & replicates | Sample size (profiles) |
|---|---|---|---|---|---|---|---|---|
| Group 1 | The trade-off between time window and sampling frequency | 24 | Uniform Distribution (1-6) | Uniform Distribution (0-24) | Periodic: cosine, cosine 2, cosine peak<br>Non-periodic: flat | N(0, 1) | 4 h/1 day X 1 & 8 h/2 days X 1<br>3 h/1 day X 1 & 6 h/2 days X 1<br>2 h/1 day X 1 & 4 h/2 days X 1 | 12,000 |
| Group 2 | The trade-off between time window and replicates | 24 | Uniform Distribution (1-6) | Uniform Distribution (0-24) | Periodic: cosine, cosine 2, cosine peak<br>Non-periodic: flat | N(0, 1) | 4 h/1 day X 1 & 8 h/1 day X 2<br>3 h/1 day X 1 & 6 h/1 day X 2<br>2 h/1 day X 1 & 4 h/1 day X 2 | 12,000 |
| Group 3 | stationary, non-stationary and asymmetric curves | 24 | Uniform Distribution (1-6) | Uniform Distribution (0-24) | Periodic:<br>Stationary (cosine, cosine 2, cosine peak)<br>Nonstationary (cosine damp, trend exp, trend linear)<br>Asymmetric (Saw-tooth)<br>Non-periodic: flat, linear | N(0, 1) | 4 h/ 1 day X 1<br>3 h/ 1 day X 1<br>2 h/ 1 day X 1 | 12,000 |
| Group 4 | Signal to Noise Ratio (SNR) 0.5:1, 1:1, 2:1, 3:1 | 24 | sqrt(2*SNR) | Uniform Distribution (0-24) | Periodic: cosine<br>Non-periodic: flat | N(0, 1) | 4 h/ 1 day X 1<br>3 h/ 1 day X 1<br>2 h/ 1 day X 1 | 12,000 |
| Group 5 | Uneven sampling (1, 2, or 4 randomly selected timepoints removed) | 24 | Uniform Distribution (1-6) | Uniform Distribution (0-24) | Periodic: cosine, cosine 2, cosine peak<br>Non-periodic: flat | N(0, 1) | 4 h/ 1 day X 1<br>3 h/ 1 day X 1<br>2 h/ 1 day X 1 | 12,000 |
| Group 6 | Missing value (1%, 5%, 10% genes missing) | 24 | Uniform Distribution (1-6) | Uniform Distribution (0-24) | Periodic: cosine, cosine 2, cosine peak<br>Non-periodic: flat | N(0, 1) | 4 h/ 1 day X 1<br>3 h/ 1 day X 1<br>2 h/ 1 day X 1 | 12,000 |

Supplementary Table 5

| Methods | 1 h/2 days | 2 h/2 days | 4 h/2 days |
|---|---|---|---|
| LS | 13s | 11s | 11s |
| ARS | 18s | 13s | 21s |
| JTK | 30s | 5s | 2s |
| RAIN | 729s | 26s | 4s |
| eJTK | 98s | 49s | 29s |
| MC | 61s | 29s | 34s |
| BC | 16s | 8s | 7s |

Supplementary Table 6

| Method | 5K | 10K | 50K | 100K | 500K |
|---|---|---|---|---|---|
| **LS** | 0.5667 | 0.5599 | 0.6785 | 0.7467 | 0.8188 |
| **JTK** | 0.6329 | 0.6141 | 0.6958 | 0.7604 | 0.7934 |
| **RAIN** | 0.6220 | 0.6781 | 0.8243 | 0.8501 | 0.8666 |
| **eJTK** | 0.6388 | 0.6008 | 0.6880 | 0.7775 | 0.8410 |
| **MC** | 0.6068 | 0.5982 | 0.7045 | 0.7514 | 0.8085 |
| **BC** | 0.4578 | 0.4947 | 0.6525 | 0.7463 | 0.7998 |

Supplementary Table 7

(A)

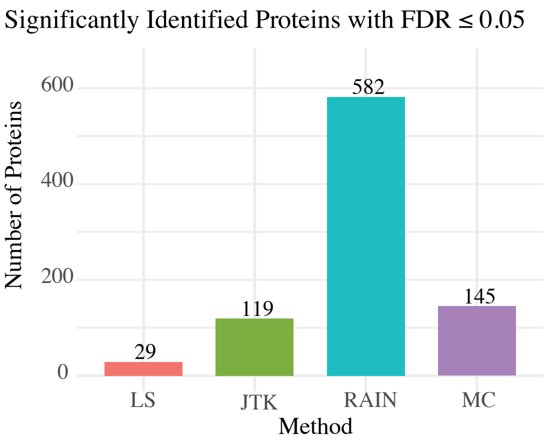


(B)

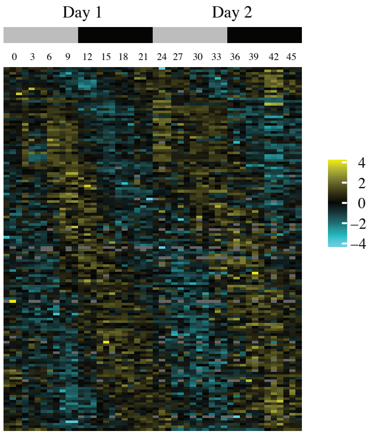


Supplementary Figure 1

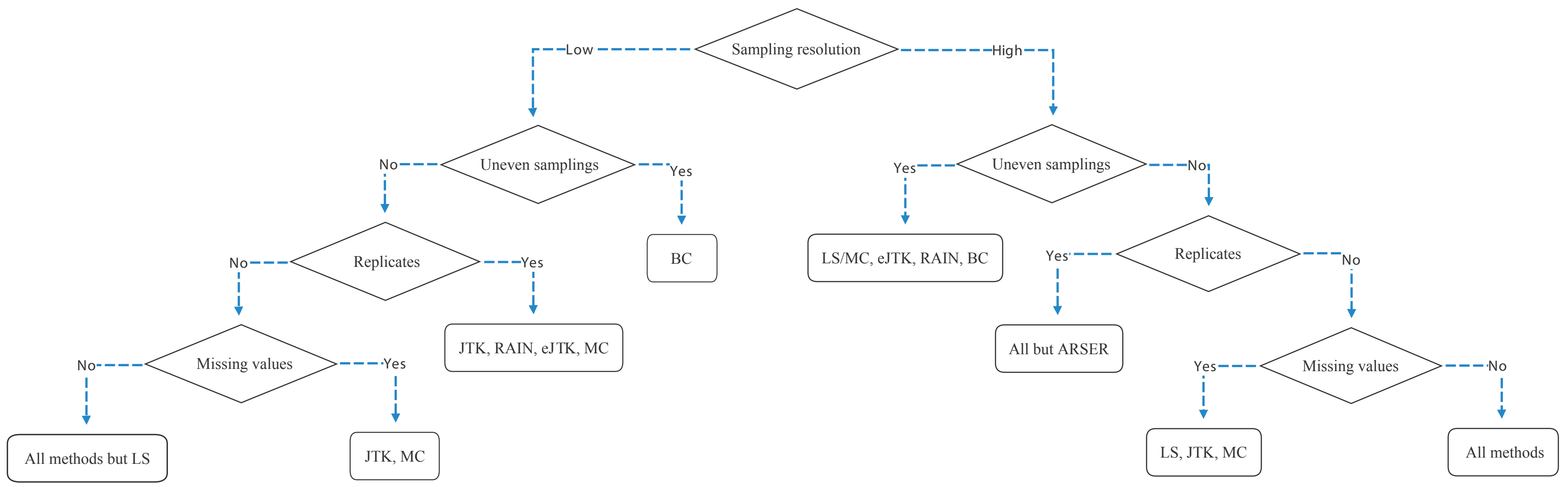


Supplementary Figure 2
